# Quantification and selection of ictogenic zones in epilepsy surgery

**DOI:** 10.1101/602490

**Authors:** Petroula Laiou, Eleftherios Avramidis, Marinho A. Lopes, Eugenio Abela, Michael Müller, Ozgur E. Akman, Mark P. Richardson, Christian Rummel, Kaspar Schindler, Marc Goodfellow

## Abstract

Network models of brain dynamics provide valuable insight into the healthy functioning of the brain and how this breaks down in disease. A pertinent example is the use of network models to understand seizure generation (ictogenesis) in epilepsy. Recently, computational models have emerged to aid our understanding of seizures and to predict the outcome of surgical perturbations to brain networks. Such approaches provide the opportunity to quantify the effect of removing regions of tissue from brain networks and thereby search for the optimal resection strategy.

Here, we use computational models to elucidate how sets of nodes contribute to the ictogenicity of networks. In small networks we fully elucidate the ictogenicity of all possible sets of nodes and demonstrate that the distribution of ictogenicity across sets depends on network topology. However, the full elucidation is a combinatorial problem that becomes intractable for large networks. Therefore, we develop a global optimisation approach to search for minimal sets of nodes that contribute significantly to ictogenesis. We demonstrate the potential applicability of these methods in practice by identifying optimal sets of nodes to resect in networks derived from 20 individuals who underwent resective surgery for epilepsy.

## 1. Introduction

Approximately 30% of people with epilepsy have refractory seizures, i.e. their seizures cannot be controlled by medication (Brodie et al., 2013, Chen et al., 2018). In these cases, the surgical removal or disconnection of the putative “epileptogenic zone” (EZ), i.e. the region of brain tissue thought to be indispensable for the generation of seizures, is a potential therapeutic option that can alleviate seizures in many cases (Rosenow and Lüders 2001, Nowell et al., 2014). The epileptogenic zone is currently defined retrospectively: diverse information is integrated by clinical teams to define targets for resection, and if seizure freedom is achieved after surgery, the EZ is assumed to have been removed (Rosenow and Lüders 2001). Unfortunately, post-operative seizure freedom rates are currently sub-optimal and not everyone who could potentially benefit from surgery is identified as a candidate (De Tisi et al., 2011, Engel et al., 2012, Fois et al., 2015, Baud et al., 2018, Engel 2018). In order to improve the success of epilepsy surgery and widen its potential usage, a better understanding of the mechanisms of seizure generation is required, and improved, quantitative methods to prospectively map the EZ need to be developed (Rummel et al., 2015, Goodfellow et al., 2016).

Computational studies of seizure generation in large-scale brain networks with the aim to inform epilepsy surgery have recently begun to emerge (Goodfellow et al., 2016, Sinha et al., 2017, Jirsa et al., 2017, Steimer et al., 2017). In an early such study we introduced a quantitative framework for prospectively evaluating the effect that surgical removal of tissue would have (Goodfellow et al., 2016). The framework proceeds by first mapping an ictogenic network i.e. a set of brain regions, together with connections between them that are important for the generation of seizures. A dynamic model is then applied to this network in order to simulate epileptiform dynamics and thereby quantify the seizure generating capability of the network. This is captured in a quantity called Brain Network Ictogenicity *(BNI)* that can be measured from simulations of the model (Petkov et al., 2014). Crucially, measuring the *BNI* for a network provides a baseline against which the effect of perturbations, such as the removal of nodes (which is a proxy for surgery), can be quantified. We recently showed that our model could accurately delineate surgeries that resulted in seizure freedom from those that did not (Goodfellow et al., 2016). Subsequent studies have added evidence for the potential use of models to guide epilepsy surgery (Sinha et al., 2017, Jirsa et al., 2017). Using models to quantify ictogenicity opens up avenues for the improvement of pre-surgical mapping. For example, putative resections can be quantitatively compared in silico, and resections providing large reductions in ictogenicity (i.e. substantial reduction in seizure occurrence) can be sought. This in itself leads to a re-imagining of the concept of the epileptogenic zone for pre-surgical planning: rather than searching for correlates of a region of tissue that should render a person seizure free, we can quantify the seizure reducing capabilities of removing many alternative sets of nodes in a network prospectively, *in silico.*

Here, we use a quantity called Set Ictogenicity *(SI)* (previously introduced as Δ*BNI* (Goodfellow et al., 2016)) to represent the extent to which ictogenicity is reduced when a given set of nodes is removed from a network. *SI* can be quantified for any potential set of nodes in a resection using the framework described above. Once the effect of removing a set of nodes can be quantified, one can compare the effect of removing different sets, and choose the set that would yield the largest reduction in ictogenicity (i.e. find the set with largest *SI*, which would be designated as a putative EZ). However, to ensure an optimal solution is uncovered, one would have to evaluate the *SI* of all possible combinations of sets of nodes within a given network. Unfortunately, this is a combinatorial optimisation problem in which the number of sets to search through quickly becomes intractable, even for moderately sized networks (Hromkovic et al., 2001, Luque et al., 2011). Though in practice constraints on the location of resections may exist, alternative strategies to exhaustive searches have to be explored. For instance, one could use heuristics that are quick but do not guarantee finding an optimum (Pearl 1984, Branke et al., 2016). For example, previous approaches (Goodfellow et al., 2016, Lopes et al., 2017, Sinha et al., 2017) applied heuristics to attempt to identify a set of nodes that optimally reduces the network ictogenicity. However, the selection of nodes of that set was solely based on the contribution of single nodes to seizure generation. In other application areas, combinatorial problems have previously been approached using global non-deterministic search strategies, like simulated annealing (Van_Laarhoven et al., 1987), evolution strategies (Beyer et al., 2001), genetic algorithms (Hromkovic et al., 2001, Luque et al., 2011) and particle swarm optimisation (Kennedy 2010). The deployment of such approaches to brain networks would enable us to gain a deeper insight into the way that ictogenesis is distributed throughout networks and facilitate the development of optimal strategies for epilepsy surgery.

Here we use computational models to study the ictogenicity of sets of nodes within a network. We use artificial networks to explore how *SI* varies across sets of nodes within networks of different topologies and to characterise the relationship between common graph metrics and *SI.* To facilitate the search for optimal resections, we develop and validate a genetic algorithm to uncover sets of nodes that optimally reduce the ictogenicity of a network. In addition, we apply the methodology to a cohort of 20 people who underwent epilepsy surgery. Finally, we discuss the potential benefits of these approaches to both enhance our understanding of epilepsy and advance pre-surgical planning in practice.

## 2. Material and methods

### 2.1. Simulation of brain dynamics

We use a mathematical modelling framework to simulate and predict the outcome of epilepsy surgery (Goodfellow et al., 2016). The framework uses intracranial electroencephalographic (iEEG) recordings (Fig 1a) to construct functional networks (Fig 1b), where nodes are associated with electrodes and edges denote interrelations between the recorded signals. We use a surrogate corrected version of mutual information (Rummel et al., 2013, Rummel et al., 2015) that detects non-linear dependencies in excess of linear relationships to define weighted edges of the network (see Supplementary material, Text S1). We then place a mathematical model on each node (Fig 1c).

**Fig. 1.**
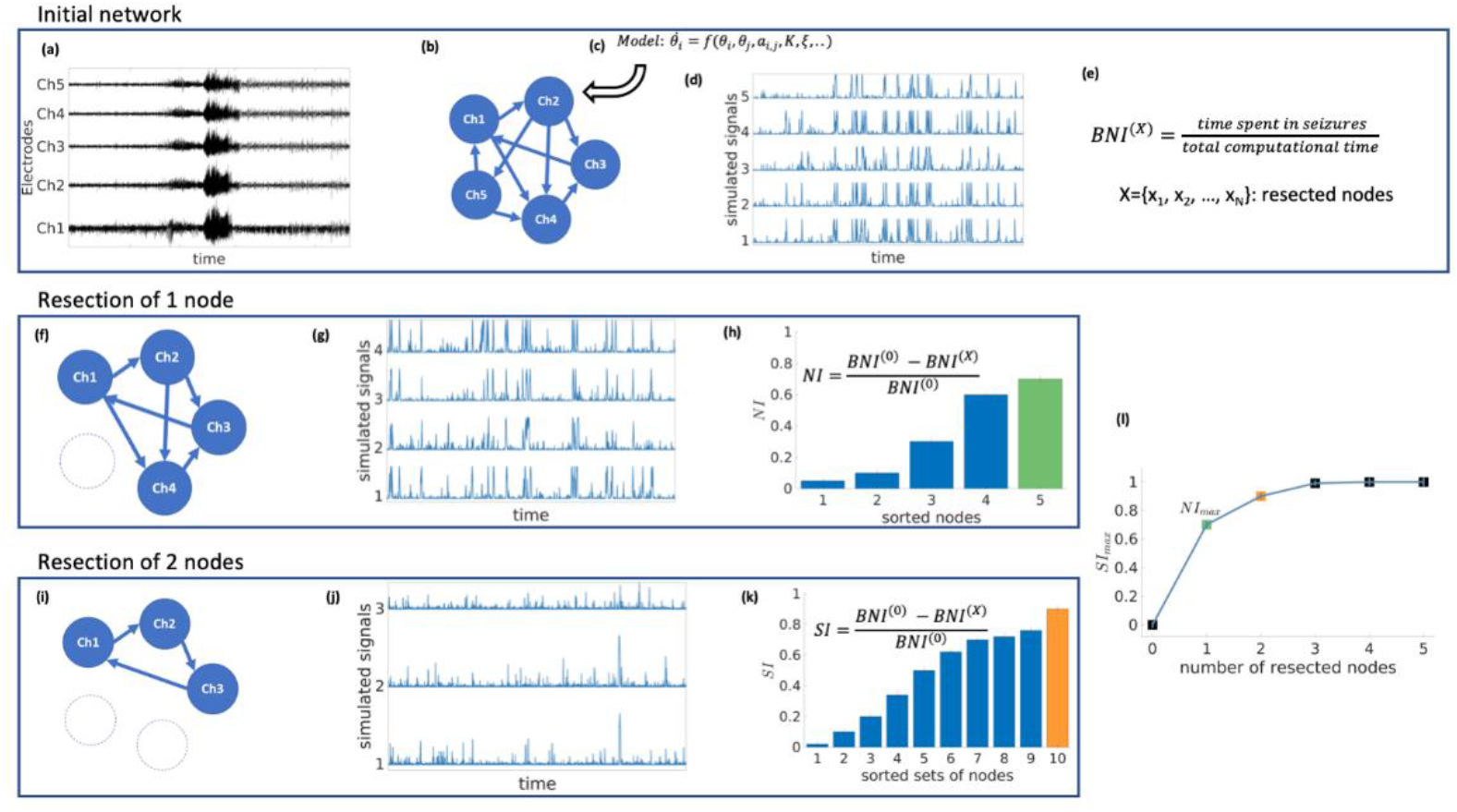
Schematic representation of the mathematical framework. From intracranial recordings (a) we construct a functional network (b). We then place a mathematical model on each node (c), simulate signals from the model (d) and calculate Brain Network Ictogenicity (e). Then perturbations are applied to the network by removing individual nodes (f) or set of nodes (i). Using the simulated signals from the perturbed networks (panels (g) and (j)), Node Ictogenicity *(NI)* and Set Ictogenicity *(SI)* are calculated for all possible combinations of resected nodes (panels (h), and (k), respectively). Finally, for every number of resected nodes the set that contributes most to the seizure generation is the one with the maximum *SI* (h, k, l). Panel (l) illustrates the *SI* of the most ictogenic sets split out by resection size.

We use the canonical theta model (Ermentrout et al., 1986, Lopes et al., 2017) which has been shown to capture relevant dynamics of neural mass models (Lopes et al., 2017); nodes can display transitions between “normal activity” (stable fixed point) and “epileptiform dynamics” (stable limit cycle) through a saddle node on invariant circle (SNIC) bifurcation. The dynamics of node *j* (*j* = 1,…, *N*) is represented by its phase *θ_j_* and obeys

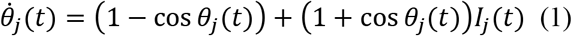

where *I_j_*(*t*) is the input current that the node receives from the other nodes in the network,

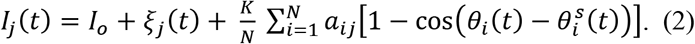

In Eq. (2) the term *I_o_* + *ξ_j_* is white uncorrelated Gaussian noise with mean *I_o_* = −1.2 and standard deviation 0.6, as used in previous studies (Lopes et al., 2017). *a_ij_* is the (*i,j*)^th^ entry of the adjacency matrix (i.e., the weighted correlation matrix of the functional network), *K* is a global scaling factor that scales the network interactions compared to the noise, and 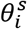 is the stable fixed point of node 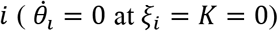. For the integration of Eq. (1) we use the Euler-Maruyama method with step size 0.01.

### 2.2. Quantification of ictogenicity

The theta model enables us to simulate brain dynamics on a network (Fig 1d). Low amplitude signals correspond to “normal activity” whilst high amplitude signals represent “epileptiform dynamics” (Lopes et al., 2017). We use the quantity Brain Network Ictogenicity *(BNI)* (Petkov et al., 2014, Goodfellow et al., 2016) to quantify the propensity of the network to generate seizures. In practice, *BNI* measures the average time that each node spends in epileptiform dynamics (*t_i_, i* = 1, …, *N*) over a sufficiently long computational time (we use 4 × 10^6^ time steps),

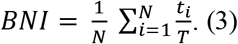

*BNI* varies between zero (all nodes display “normal activity” all the time) and one (all nodes show “seizure activity” all the time). The changes in *BNI* upon removal of individual nodes or set of nodes allow us to quantify their ictogenicity. Here, removal of nodes is implemented by setting all incoming and outgoing connections from and to other nodes to zero (Figs 1i-k). To measure the impact of removing a set of nodes on *BNI*, we define Set Ictogenicity *(SI)* as

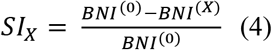

where *BNI*^(0)^ is a reference value for the unperturbed network, and *BNI*^(*X*)^ denotes the *BNI* value after the removal of all *n* nodes in the subset *X* = {*x*_1_,…, *x_n_*}. We tune the parameter *K* in the model such that *BNI*^(0)^ equals 0.5 (Goodfellow et al., 2016, Lopes et al., 2017) which means that on average the network spends half of the computational time in epileptiform dynamics. We consider this value as a useful reference because it enables us to study the result of network perturbations more efficiently (a realistic value of *BNI*^(0)^ would be much smaller, and changes of *BNI* upon node removals would be more difficult to measure). Note that for *n* = 1 in Eq. (4), *SI_X_* is equivalent to the Node Ictogenicity *(NI)*, previously introduced in Goodfellow et al., (Goodfellow et al., 2016), which measures the effect of removing a single node in *BNI* (Figs 1f-h). *SI_X_* is a succinct term for *ΔBNI*, which was introduced in Goodfellow et al., (Goodfellow et al., 2016). Larger *SI_X_* values denote greater contribution of the considered set *X* to seizure generation. *SI_X_* takes a value of one when the removed set of nodes leads to elimination of epileptiform dynamics, whilst zero or negative values denote that removing those nodes did not reduce the network ictogenicity. In this study, we set all negative *SI_X_* values to zero. More details about the calculation of *BNI* and *SI* are given in Supplementary material, Text S2.

### 2.3. Exemplar networks and topological properties

In order to explore and understand the effect of perturbations in different network structures, we first study *SI* in exemplar networks. We consider both directed and undirected artificial networks with random and “scale-free” (i.e. generated by the Barabasi-Albert algorithm (Barabasi 1999) and the static model (Goh et al., 2001)) topologies, comprising 20 and 40 nodes. We also note that in contrast to the weighted functional networks inferred from patient data (see section: Patient information and data), we used binary artificial networks for simplicity. We further consider common graph theory measures such as the degree, betweenness centrality, clustering coefficient and eigenvector centrality (Newmann 2007, Rubinov and Sporns 2010) to study how *SI* relates to these topological properties (eigenvector centrality was only computed for undirected networks, because it is undefined for directed networks).

### 2.4. Resection strategies

In a network of *N* nodes there are 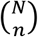 distinct subsets of size *n*. Therefore, in order to evaluate how *SI* is distributed within a network of size *N*, and to definitively identify the set with the highest *SI*, one could use an exhaustive (or brute-force) search, which would require 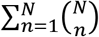 calculations. We use such an approach herein to calculate the ‘ground truth’ distribution of *SI*. For networks of size relevant for the study of the brain, the brute-force approach quickly becomes intractable. For example, a 40-node network exhaustive search would require 2^40^ – 1 ≈ 10^12^ calculations. We therefore need to develop computationally tractable methods for studying *SI*. Previous studies (Goodfellow et al., 2016, Lopes et al., 2017) have used heuristic methods based on recursively adding a single node to build up an optimal set. One method, which we refer to as ‘simple ordering’ (Goodfellow et al., 2016) calculates *NI_X_* for all possible single node removals (i.e. *N* initial calculations). Nodes are ranked according to their *NI_X_* values and added sequentially to the set. Each time a new node is added, the *SI* of the new set is calculated, with termination when *SI* is greater than 0.99. An extension to this method, which we refer to as ‘recurrent ordering’ recalculates the distribution of *NI* after the removal of each node to ensure that in every iteration the node with the highest *NI* of the perturbed network is added.

In addition to these heuristics, we test a resection strategy based on optimising *SI* with a genetic algorithm. Genetic algorithms are stochastic search methods based on mimicking natural selection, in which an evolving population of candidate problem solutions is used to find an optimal solution (Avramidis and Akman 2017, Luque 2011, Zhou et al., 2011). A typical genetic algorithm (GA) starts with a population that comprises candidate solutions (called individuals). Each individual is evaluated by a fitness (objective) function which quantifies how successfully the individual solves the problem. Based on the fitness scores, the genetic algorithm creates a new population of individuals by performing a number of stochastic genetic operations (i.e. crossover, mutation, selection), and keeps the best solutions generated (those that minimize the objective function). This process continues for multiple iterations (called generations) until convergence to an optimum solution is achieved. Multi-objective genetic algorithms (MOGAs) optimise the given problem for more than one objective function, returning Pareto optimal solution sets which represent the optimal trade-off between the objectives (Zhou et al., 2011, Avramidis and Akman 2017). Here, we use the Non-dominated Sorting Genetic Algorithm-II (NSGA-II) (Deb et al., 2002, Avramidis and Akman 2017). Particularly, we use the implementation of the algorithm included in the MATLAB Global Optimisation Toolbox (version R2017b) and follow the optimisation protocol from Avramidis et al., (Avramidis and Akman 2017). Given that our purpose is to find the smallest set with the largest *SI*, we use two objective functions that minimize the number of nodes removed as well as the quantity 1 – *SI*. After multiple generations, the algorithm returns optimal sets of nodes with different resection sizes. Due to the stochastic nature of the genetic algorithm, we execute eight independent runs in each case utilizing the convergence metrics from Avramidis et al., (Avramidis and Akman 2017 and references therein) to assess whether the algorithm is robust and reliable at identifying optimal sets. In short, the convergence metrics evaluate the spread of the optimal sets in the two-dimensional plane of the objective functions. For efficiency, we set the genetic algorithm to discard sets of nodes larger than half of the network size by attributing them arbitrarily large objective values.

All the aforementioned search methods were evaluated on 20-node networks. In 40-node networks we did not use the “ground truth” strategy, given that it is computationally intractable to calculate Set Ictogenicity for 2^40^ – 1 ≈ 10^12^ sets. We therefore introduce in this case an additional approach: a ‘random search’ heuristic which picks at random a sample of sets of nodes for every resection size and takes as a solution the set with the maximum *SI*. For comparison purposes, we consider a sample size equal to the number of sets evaluated by the genetic algorithm across all generations. The number of samples per resection size was considered proportional to the logarithm of all possible sets of the respective size. This enabled a denser sampling at small resection sizes where the optimum solution is expected to be found.

### 2.5. Comparison between resection strategies that are based on Node and Set Ictogenicity

The ground truth is the only strategy that guarantees the detection of the most ictogenic set, because it is the only one that has access to the *SI* of all sets of nodes (Figs 1h, k, l). Thus, for the 20-node networks we compare all strategies to the optimum solution found from the ground truth. In order to evaluate how close a given solution is to the highest *SI* observed in the ground truth, we compute the Δ*SI*,

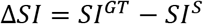

where *SI^GT^* is the highest *SI* observed in the ground truth and *SI^s^* is the *SI* of the optimum solution detected by a strategy *S* (for a given resection size). Our aim is to find a strategy that may find *SI^GT^* (Δ*SI* = 0) while avoiding the inefficient and exhaustive search performed in the ground truth.

In the case of 40-node networks, we cannot calculate the ground truth and therefore we use the solution from the genetic algorithm (*SI^GA^*) as a reference to compare with other strategies,

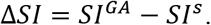

Note that in this case the Δ*SI* could be negative, because the genetic algorithm might be outperformed by another strategy.

### 2.6. Patient information and data

We analyse intracranial electroencephalographic recordings from 20 patients (15 female, 5male; median age 31 years, IQR 16 years, range 10-66 years) who underwent presurgical monitoring at Inselspital Bern. 13 of them were free from disabling seizures and auras for at least one year after surgery (Engel I), whereas the remaining 7 did not show worthwhile improvement (Engel IV). Before and after surgery, high resolution MRI images were acquired, as well as post-implantation CT images in order to identify the position of the implanted electrodes and the exact location of the resected brain tissue. Further details about this procedure can be found in Rummel et al., (Rummel et al., 2015). An experienced epileptologist/electroencephalographer (K.S.) visually inspected all the recordings and identified the onset and termination of a representative seizure as well as any channels that had to be removed from the analysis due to the presence of permanent artefacts (<5% of channels). All signals were down-sampled to a sampling rate of 512 Hz, re-referenced against the median of all the artefact-free channels and band-pass filtered (forward and backward filtering to minimize phase distortion) between 0.5 and 150 Hz using a fourth-order Butterworth filter. All the patients gave written informed consent that their imaging and EEG data might be used for research purposes, and retrospective data analysis has been approved by the ethics committee of the Canton of Bern/Switzerland.

## 3. Results

The results section is arranged as follows. In order to better understand how sets of nodes within networks contribute to ictogenicity, we first study how *SI* is distributed across subnetworks of different sizes in artificially generated scale-free and random networks. We subsequently assess the relationship between *SI* and common graph theory metrics and test to what extent these metrics can predict the optimal set. We then apply a genetic algorithm and heuristics to the problem of finding the optimal set before finally testing the genetic algorithm on patient data.

### 3.1 Set Ictogenicity in different network topologies

In a network of size *N*, for a given subset (collection) of *n* nodes, *X* = {*x*_1_, …, *x_n_*}, where *n* < *N*, we use *SI_X_* to quantify the reduction in ictogenicity that is achieved by removing all nodes in *X* from the network. To gain insight into how seizures arise in networks, we first seek to uncover what the relationship is between the obtained reduction in ictogenicity (*SI_X_*) and the number of nodes that are removed (*n*), and how this depends on network topology. For computational tractability we initially study 20-node artificially generated networks, considering the removal of up to ten nodes (i.e. up to half of the network). This network size is tractable for analysis using a brute-force approach and is relevant in the clinical context, where iEEG implantation schemes for some people may comprise around 20 electrodes, and investigations of standard clinical scalp EEG typically yields 19 channels.

Figs 2a, b, e and f demonstrate how *SI* is distributed in exemplar directed and undirected networks with scale-free and random topologies. We observe that the variance in *SI* is larger in the directed scale-free networks (Fig 2a) compared to random (Figs 2b and f) and undirected scale-free (Fig 2e) networks. Fig 2a shows that in the directed scale-free networks we studied, *SI* can take values between zero and one, depending on the set that is removed. Approximately 10% of sets do not reduce ictogenicity when removed. A further 20% completely eliminates epileptiform dynamics when removed, but the effect of the remaining sets is distributed approximately uniformly across *SI* values. In contrast, Figs 2b, e and f demonstrate that the *SI* distribution of random and scale-free undirected networks is more concentrated at high values, with very few sets having no effect on ictogenicity.

**Fig. 2.**
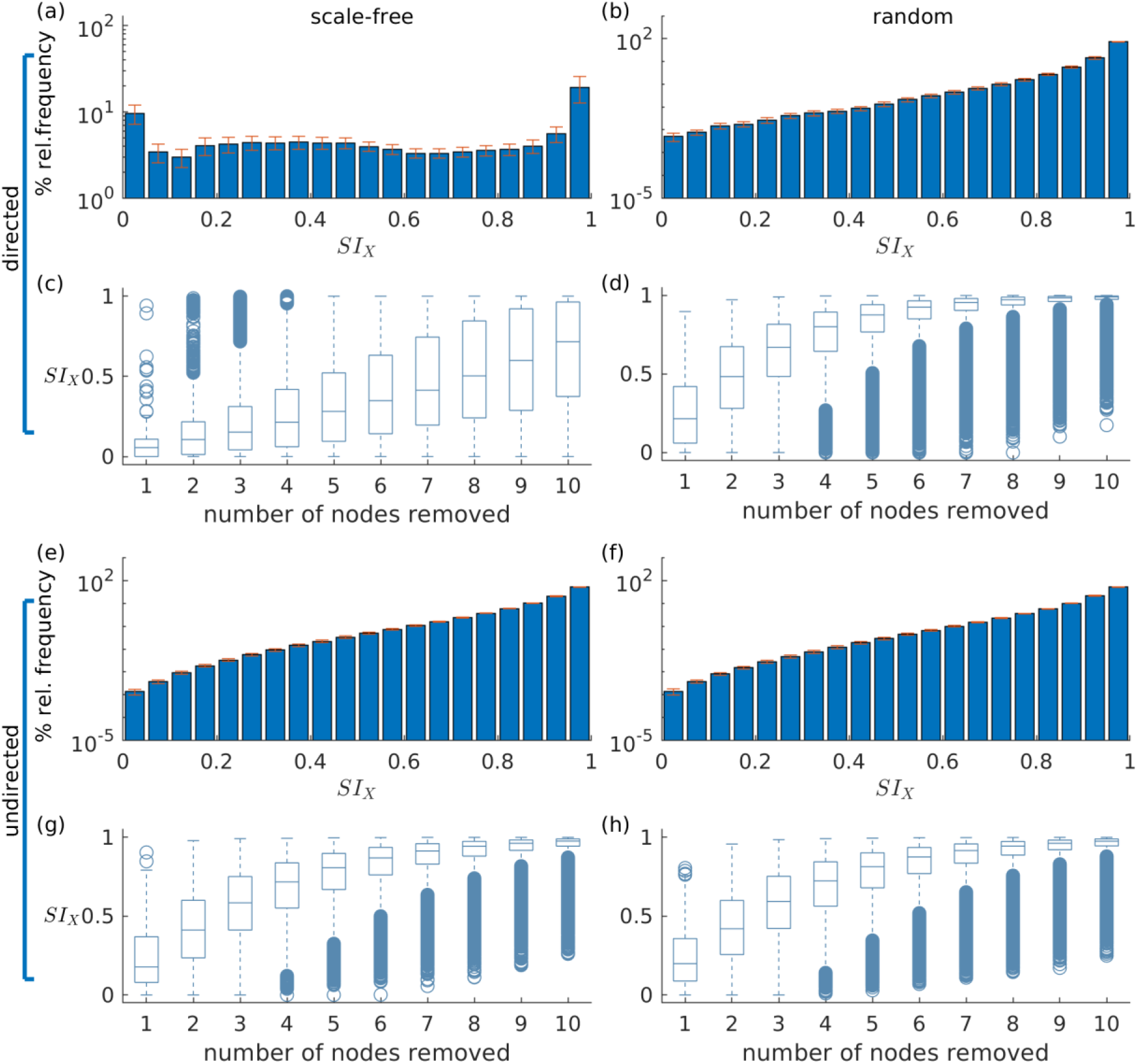
The distribution of Set Ictogenicity *(SI)* depends on network topology and resection size. Panels (a) and (b) display the average *SI* distribution for ten directed artificial scale-free (a) and random (b) networks, respectively. Error bars denote the standard error across the ten network realizations. Panels (c) and (d) show *SI* distributions split out by resection size (i.e. number of nodes removed) for the networks of panels (a) and (b), respectively. Panels (e)-(h) are similar to panels (a)-(d), but for undirected networks. Parameters: network size *N* = 20; in the directed networks the in and out degree is 2, whilst in the undirected the mean degree is 2; scale-free degree distribution exponent *γ* = 3.

In Figs 2c, d, g and h *SI* is broken down by resection size. In all networks studied, the average *SI* increases as the size of the resected set increases. However, in the scale-free directed networks (Fig 2c) we studied, the relationship between resection size and average *SI* is approximately linear, and in random (Figs 2d and h) and undirected scale-free networks (Fig 2h), large average *SI* values are reached more readily for small set (resection) sizes. Furthermore, whilst in the directed scale-free networks we studied there are few sets with very large *SI*, in random and undirected scale-free networks there are instead few low *SI* values. We observe that typically, small resections have on average greater impact (i.e. higher *SI)* in random networks, compared to equivalent resections in directed scale-free networks.

### 3.2. Set Ictogenicity and graph theory measures

Fig 2 demonstrated that *SI* depends on network topology. To further understand this relationship, we investigated to what extent *SI* is related to common graph theoretic properties of nodes inside the sets selected for removal. We thus computed the Spearman’s rank correlation between *SI* and the average degree, average betweenness centrality and average clustering coefficient of removed sets in directed scale-free and random networks. Fig 3 shows that *SI* is correlated with average degree and average betweenness centrality (median correlation larger than *0.6* for most resection sizes, see Figs 3a and b), but not with the average clustering coefficient in both scale-free and random networks (directed and undirected, see Fig S3 for the latter). Analysing different resection sizes we see that there are differences in correlation between directed scale-free networks and the other topologies we studied. In random networks, for example, the correlation between *SI* and average degree and betweenness centrality is high for small resection sizes but decreases for larger resections (Figs 3d and e). In contrast, correlation between degree and *SI* increases with resection size for directed scale-free networks (Fig 3a) and is relatively flat for betweenness centrality (Fig 3b). The low correlation for large resection sizes in random networks, particularly for betweenness centrality, is likely to be a consequence of the fact that most large sets have the same *SI* in random networks, as found in Fig 2d. Therefore, *SI* would be independent from measures of the constituent nodes.

**Fig. 3.**
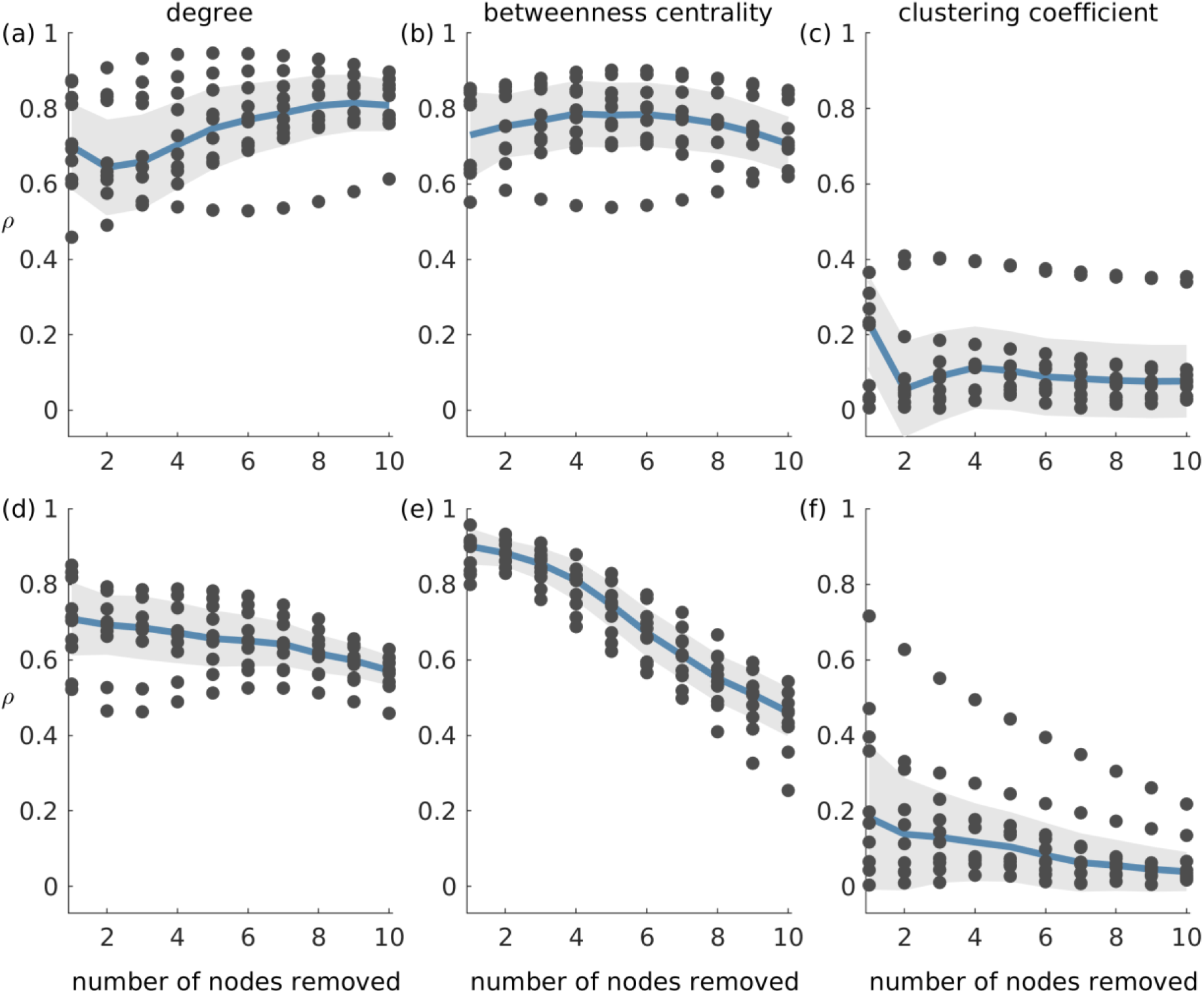
Absolute Spearman’s correlation (*ρ*) between *SI* and average graph theory measures of the nodes in removed sets. Panels (a)-(c) correspond to scale-free directed networks while panels (d)-(f) represent random directed networks. Each column shows *ρ* between *SI* and a different network measure: (a) and (d) average degree; (b) and (e) average betweenness centrality; and (c) and (f) average clustering coefficient of removed nodes. Ten network realizations were considered per network topology, hence the 10 dots for each resection size (i.e. number of nodes removed). The blue line represents the median across the network realizations and the shaded area displays the median absolute deviation. The parameters were the same as in Fig 2.

### 3.3. Identifying optimal resections

Having explored properties of *SI* distributions, we now turn to the problem of finding the set that optimally reduces ictogenicity upon removal, which is analogous to identifying the optimal resection in epilepsy surgery. Fig 3 demonstrated that the average degree and average betweenness centrality of removed sets are correlated with *SI.* This implies that these graph theory measures may potentially be used to find the sets with highest *SI.* We sought to test this by asking whether sets that produce maximum reductions in average degree or average betweenness centrality also produce maximal *SI* values. In order to identify maximal *SI* values, we calculated *SI* for all possible subsets of artificial networks with 20 nodes, which we refer to as the ground truth. Note that for a given network and resection size, there may exist multiple sets that produce maximal reductions in average degree or average betweenness centrality. We therefore calculated the average difference between the maximum *SI* (ground truth) and the *SI* of each set that yields maximum reduction average degree and betweenness centrality (Δ*SI*). We henceforth denote sets that yield maximal reduction in average degree and average betweenness centrality as *SI_deg_* and *SI_bet_*, respectively.

Fig 4 shows the results of this analysis for directed scale-free (Figs 4a and c) and random (Figs 4b and d) networks. We observe that in both scale-free and random directed networks, sets corresponding to *SI_deg_* and *SI_bet_* can yield *SI* values close to the optimal *SI* as defined by the ground truth (i.e. Δ*SI* = 0). In addition, the *SI_deg_* and *SI_bet_* sets always have lower Δ*SI* than the average *SI* from sets of the same size, which indicates that the reduction in average degree and average betweenness centrality are useful ways to find optimal sets (see the dashed line in the figure which represents the average Δ*SI* of every possible set of nodes). We also find that Δ*SI* gets smaller with increasing resection sizes, which is a consequence of how the distribution of *SI* changes with the resection size (i.e. as the number of nodes removed becomes large, *SI* becomes large in general, see Fig 2). However, we also observe that sets producing the same reduction in average degree or average betweenness centrality may have different *SI* values, as shown by the large error bars in Figs 4a and b. This means that information regarding the degree or centrality alone is insufficient to identify the actual optimal set, since we would need further information to identify which of the sets corresponding to *SI_deg_* and *SI_bet_* would have highest *SI.* Furthermore, Δ*SI* may be quite large for some realizations of directed scale-free networks (see the existence of outliers in Figs 4a and b). In contrast, in random directed networks average degree and average betweenness centrality more accurately identify sets with optimal *SI* (see Fig 4c and d), particularly at larger resections (which is to be expected given that most sets yield *SI* close to the highest at large resection sizes independent of their constituent nodes). Results for undirected networks are shown in Fig S4 where we also explored whether sets that produce maximum reduction in eigenvector centrality also produce maximal *SI* values. We also performed this analysis for clustering coefficient and found found that the corresponding Δ*SI* values were very large which means that clustering coefficient was not able to identify the optimal set as defined by the ground truth.

**Fig. 4.**
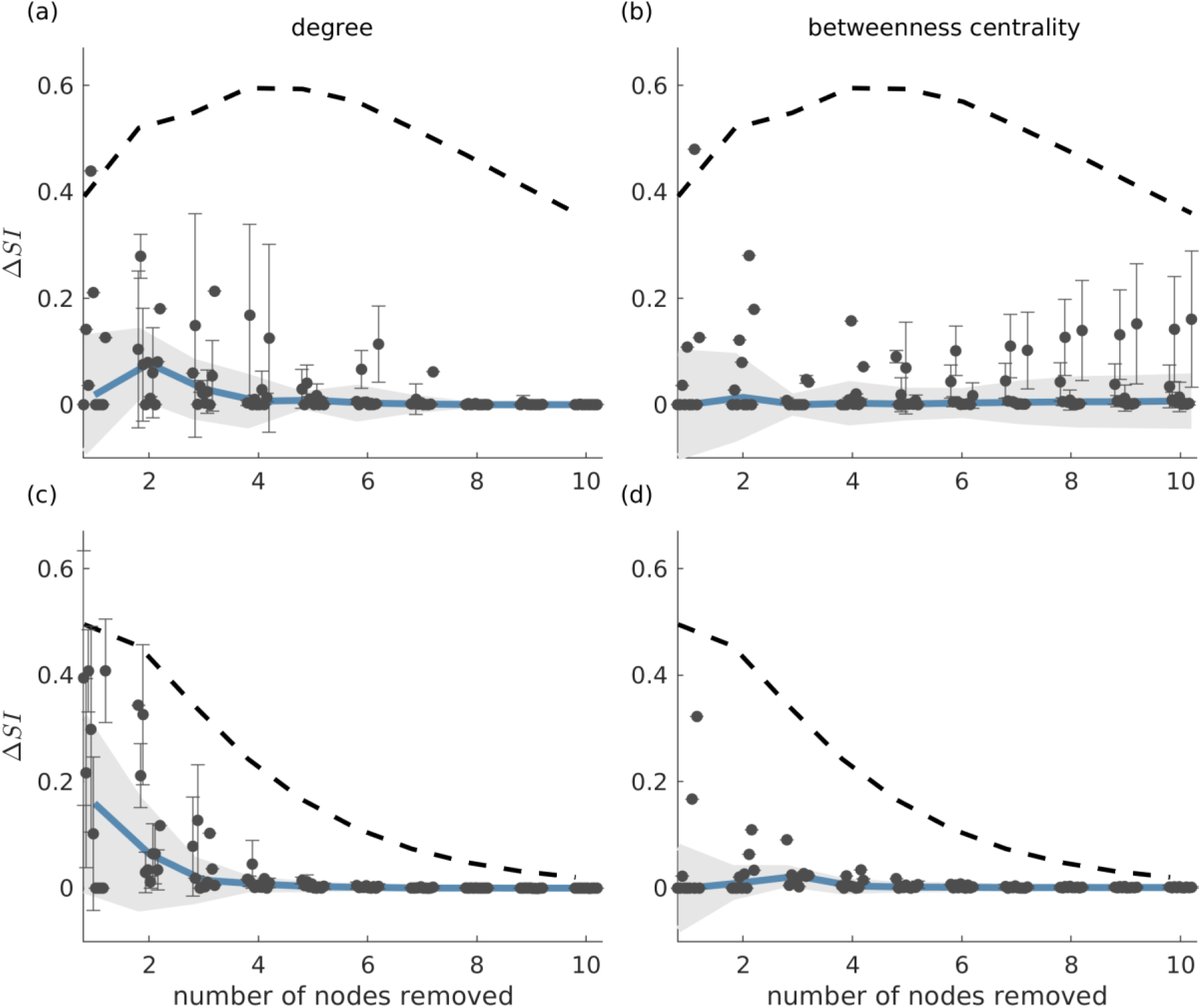
Difference between the *SI* value of the most ictogenic set as identified from the ground truth and the average *SI* of the sets which caused a maximal reduction in average degree (panels (a) and (c)), and average betweenness centrality (panels (b) and (d)), i.e. Δ*SI*, as a function of resection size. Error bars denote the standard deviation of the Δ*SI* values across the different sets that yield the maximal reduction in average degree or betweenness centrality when removed. The blue curve describes the median of the Δ*SI* values across 10 network realizations (black dots) and the shaded area their median absolute deviation (the dots are slightly shifted in the x-axis for better visualization). Panels (a) and (b) correspond to scale-free directed networks, whilst panels (c) and (d) to random directed networks. The dashed line denotes the average Δ*SI* between the *SI* of the ground truth most ictogenic set and the *SI* of all other possible sets (also averaged over the 10 network realizations). Parameters are the same as in Fig 2.

### 3.4. Alternative strategies for the identification of the most ictogenic sets

Fig 4 showed that graph theory measures may often be used to identify the most ictogenic set. However, they may not be reliable for certain network realizations, particularly in directed scale-free networks. Therefore, to find the set of nodes with maximal ictogenicity we should calculate *SI.* However, calculating the *SI* for all possible sets is challenging computationally due to the large number of possible sets, particularly in large networks. Therefore, in order to study larger networks, and to find optimal resections in general for practical applications, efficient methods are required to find sets with optimal *SI.* Here, we study two previously used heuristics, along with the NSGA-II genetic algorithm (Deb et al., 2002, Avramidis and Akman 2017). The heuristics we use are the simple ordering and recurrent ordering methods, which are based on the contribution of individual nodes to seizure generation. In contrast to building pseudo-optimal sets recursively, the genetic algorithm makes stochastic searches in the space of all possible sets of nodes using natural selection criteria.

In Fig 5 we compare these methods against the ground truth in 20-node directed networks. We find that whilst both simple and recurrent ordering are able to identify solutions close to the highest *SI*, the genetic algorithm is the only approach that uncovers the optimal solution in all cases (the red line overlaps with the green line in Fig 5c, and Δ*SI* = 0 in panels (f) and (i)). We further observe that in general the recurrent ordering provides better solutions than the simple ordering. These strategies performed differently for different network topologies, with scale-free directed networks being less amenable to the heuristic approaches than random networks (see the higher Δ*SI* values in panels (d) and (e) compared to (g) and (h)). In undirected networks we observe similar results, though in this case both the genetic algorithm and the recurrent ordering strategy are able to find the sets with highest *SI* (see Fig S5). We note that the genetic algorithm and the recurrent ordering approaches perform better than the heuristics based on graph theory measures (compare, for example, Figs 5e and h with Fig 4), further motivating the benefit of calculating *SI.* For all considered network topologies of this study, we ensured that the genetic algorithm converged across multiple independent realizations. The *SI* values of the optimal sets across multiple runs as well as the convergence metrics for exemplar networks can be found in Supplementary material (Figs S1 and S2).

**Fig. 5.**
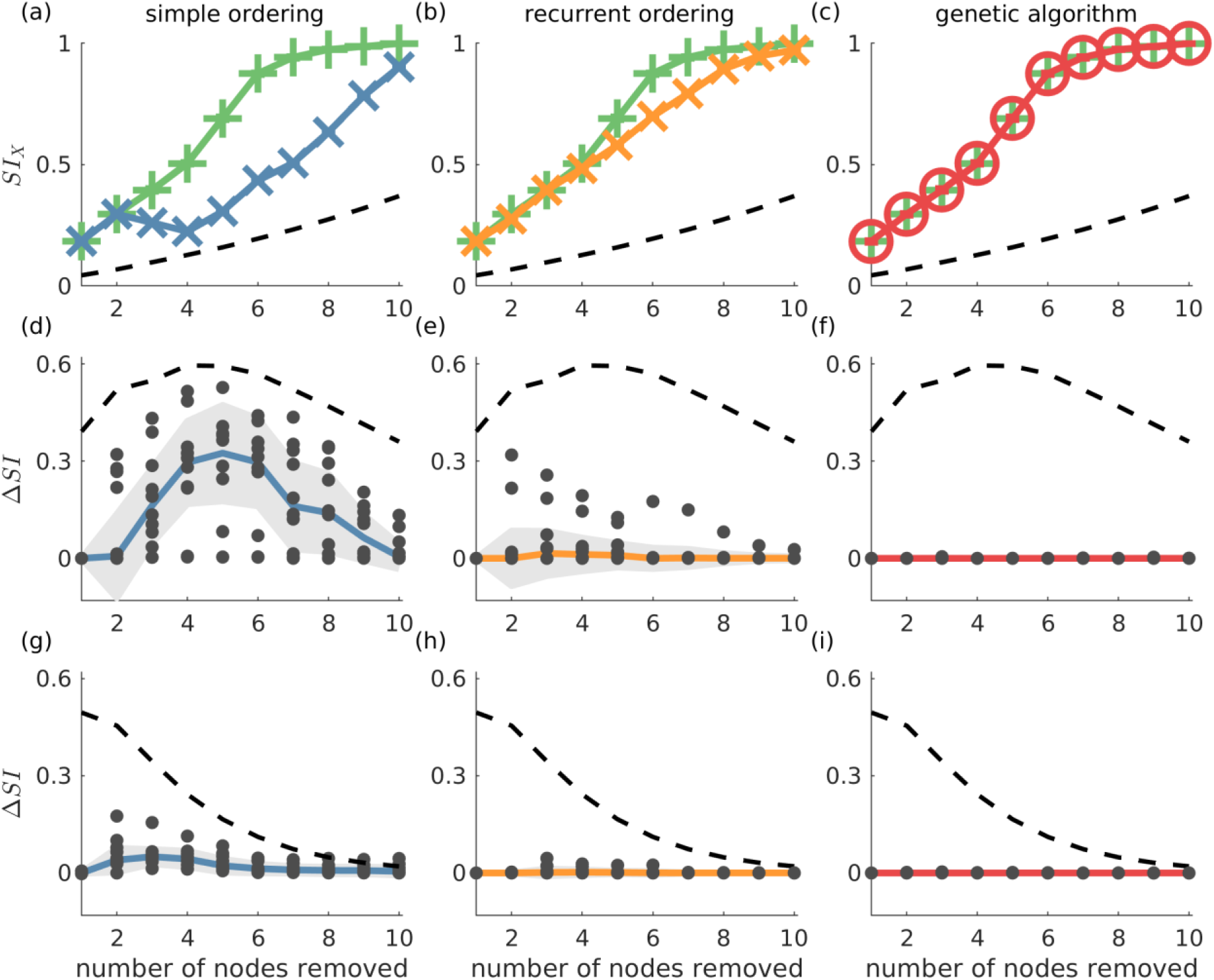
The genetic algorithm is the only resection strategy that identifies the most ictogenic set in all considered networks. Ground truth is compared with simple ordering (first column), recurrent ordering (second column) and the genetic algorithm (third column). Panels (a)-(c) display *SI* values over the number of resected nodes in an exemplar artificial 20-node scale-free directed network, where the green lines represent the ground truth (i.e. the most ictogenic sets), whilst the blue, orange, and red lines display the *SI* of the sets found by each of the search strategies. The second and third rows show the Δ*SI* between the *SI* value of the ground truth and the set identified by each search strategy in scale-free and random directed networks, respectively. The dashed line represents the average *SI* (or Δ*SI*) of all possible combinations of nodes for each resection size and serve as a reference for comparison. The dots in panels (d)-(i) correspond to Δ*SI* values obtained for 10 different network realizations, the solid lines depict the median of Δ*SI* values, and the shaded area represents the median absolute deviation. Parameters were the same as in Fig 2. Additionally, the genetic algorithm was run for 100 generations with a population size of 200.

The analysis above showed that the genetic algorithm can identify sets with the largest *SI* in all of the 20-node networks considered. In addition, the recurrent ordering strategy also found solutions relatively close to the optimum *SI* (see Fig 5e and h). Therefore, we sought to explore whether these optimisation strategies may also achieve good performances on larger 40-node networks. For these larger networks, we restricted our analysis to directed scale-free networks given that they proved to be the more difficult to approach in our analysis of 20-node networks. As explained in Methods, we do not compute the ground truth in 40-node networks, since the number of all possible sets of nodes is too large. Instead, we use the genetic algorithm as a proxy for the ground truth and compare this to the simple and recurrent ordering heuristics. In addition, we employed a random search heuristic that searches through a solution space whose size is equal to the one of the genetic algorithm in order to test the uplift in performance of the latter compared to a stratified random approach.

Fig 6 shows that in the 40-node scale-free directed networks we studied, the genetic algorithm clearly outperforms all the other strategies. Note that in this figure Δ*SI* > 0 means that the genetic algorithm finds solutions with larger *SI* than the other approaches. We find that the uplift in performance of the genetic algorithm has a maximum at around sets of size 10 and then decreases for larger sets. This is due to larger resections being more likely to have *SI* = 1, as observed in our study of 20-node networks. This is further supported by the V-shaped dashed guideline curves corresponding to random removals which decrease for resection sizes larger than 15. In contrast, in undirected networks all approaches detect similar *SI* solutions (see Fig S6).

**Fig 6.**
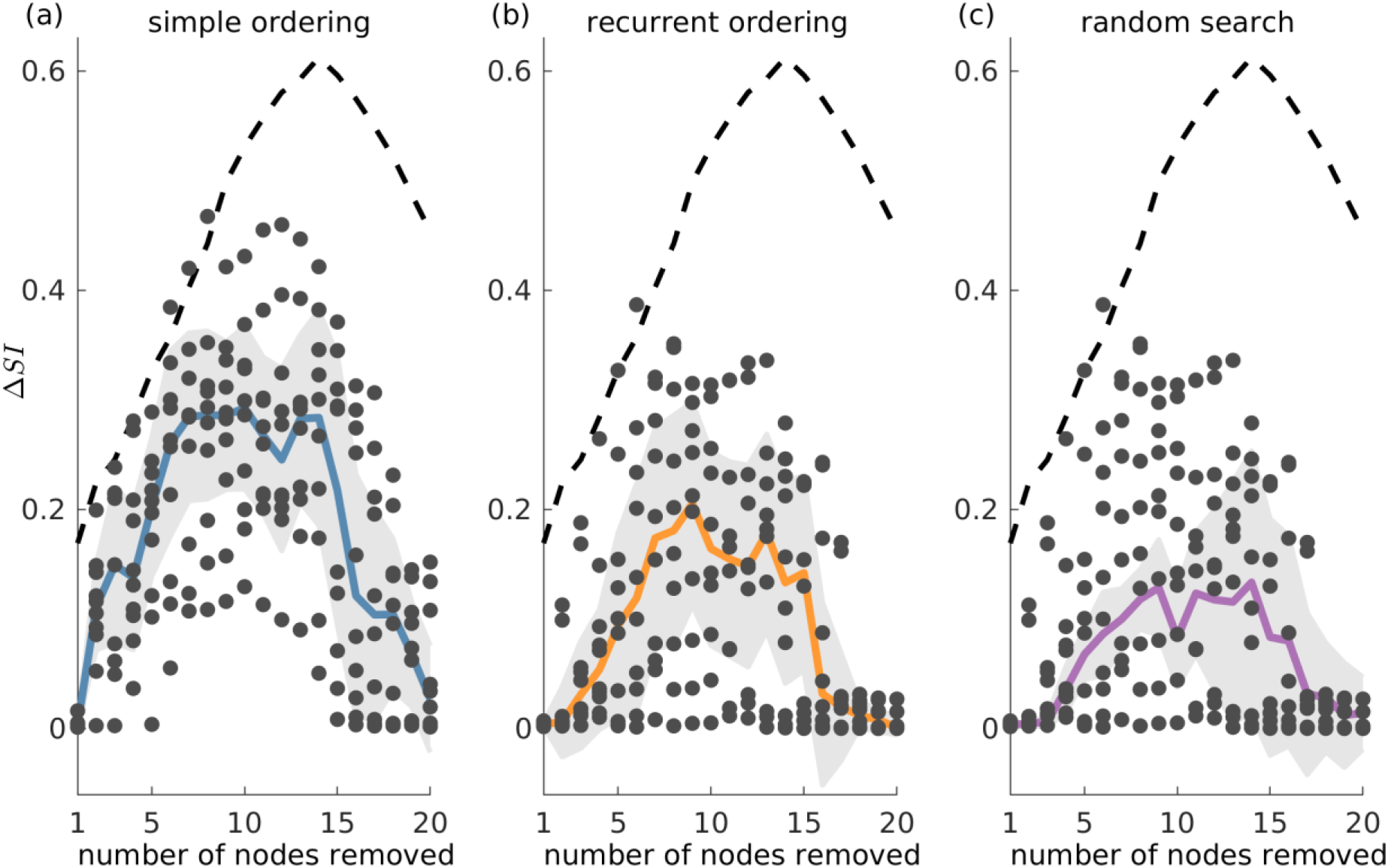
The genetic algorithm outperforms both simple and recurrent ordering as well as the random search heuristic in 40-node scale-free directed networks. Δ*SI* denotes the difference between the *SI* value of the optimal set as detected by the genetic algorithm and the *SI* solution found by (a) simple ordering, (b) recurrent ordering, and (c) random search across different resection sizes. The solid lines depict the median of the dots which correspond to the Δ*SI* values across 10 network realisations, whilst the shaded area illustrates the median absolute deviation. The dashed line represents the difference between the optimal set as detected by the genetic algorithm and the average of 20,000 random sets for each resection size (for resections up to three nodes, we considered all possible sets, since there were fewer than 20,000). Parameters: mean in and out degree equal 4; population size 200, number of generations equal 150.

### 3.5. Identifying optimal resections in patient data

Our analysis of artificial networks demonstrates that the genetic algorithm is a good strategy for identifying sets of nodes with the largest *SI*. We therefore sought to test the application of the genetic algorithm to networks inferred from a cohort of 20 people with pharmacoresistant epilepsy who underwent resective surgery (see Material and methods). Using iEEG recordings, we constructed a functional network for each patient (see Supplementary material, Text S1) and used the genetic algorithm to identify the most ictogenic sets. Here, the network nodes correspond to iEEG channels, which in turn represent the brain tissue in the vicinity of the electrodes. Following (Goodfellow at al. 2016) we defined the optimal set as the smallest set for which the *SI* exceeds 0.99. Note that as described in the Methods section, we executed multiple independent runs of the genetic algorithm, which is inherently stochastic, in order to obtain robust results. We observed that across the independent realizations we could obtain multiple optimal sets for a given patient (i.e. different sets of nodes with the same size and *SI* value).

In order to test the validity of predictions of the model, we compared them to the actual resections the patients underwent, and whether they were rendered seizure free as a result. Since the algorithm yielded multiple potential resections we calculated the overlap between the actual resected tissue and each of the optimal sets. Fig 7 shows the largest overlap per patient, with individuals grouped by postsurgical outcome. We found significantly larger overlaps for individuals who had good postsurgical outcome (Engel I) compared to those who had poor outcome (Engel IV) (see Fig 7a). In addition, we found that three out of the seven patients with poor post-surgery outcome presented zero overlap, meaning that our methods suggested completely different resections compared to those that were performed. We also compared the actual resected tissue with equal size random sets of nodes and found that in all cases the overlap with the random sets was lower than that predicted by our methods. The only exceptions were again the three cases with zero overlap with our predictions. Using the overlap as a classifier we found a sensitivity of 0.92, a specificity of 0.71, and an area under the curve (AUC) of 0.87 (see Fig 7b), which suggests our methods are reliable for classifying into outcome classes at the individual level. Interestingly, using graph theory measures alone Engel class I and IV patients could not be separated (Fig S7). However, we found that if *SI* and the genetic algorithm are used to calculate the optimal size of the resection as a first step, eigenvector centrality and strength were able to separate the two groups well, whereas betweenness centrality could not (Fig S8).

**Fig 7.**
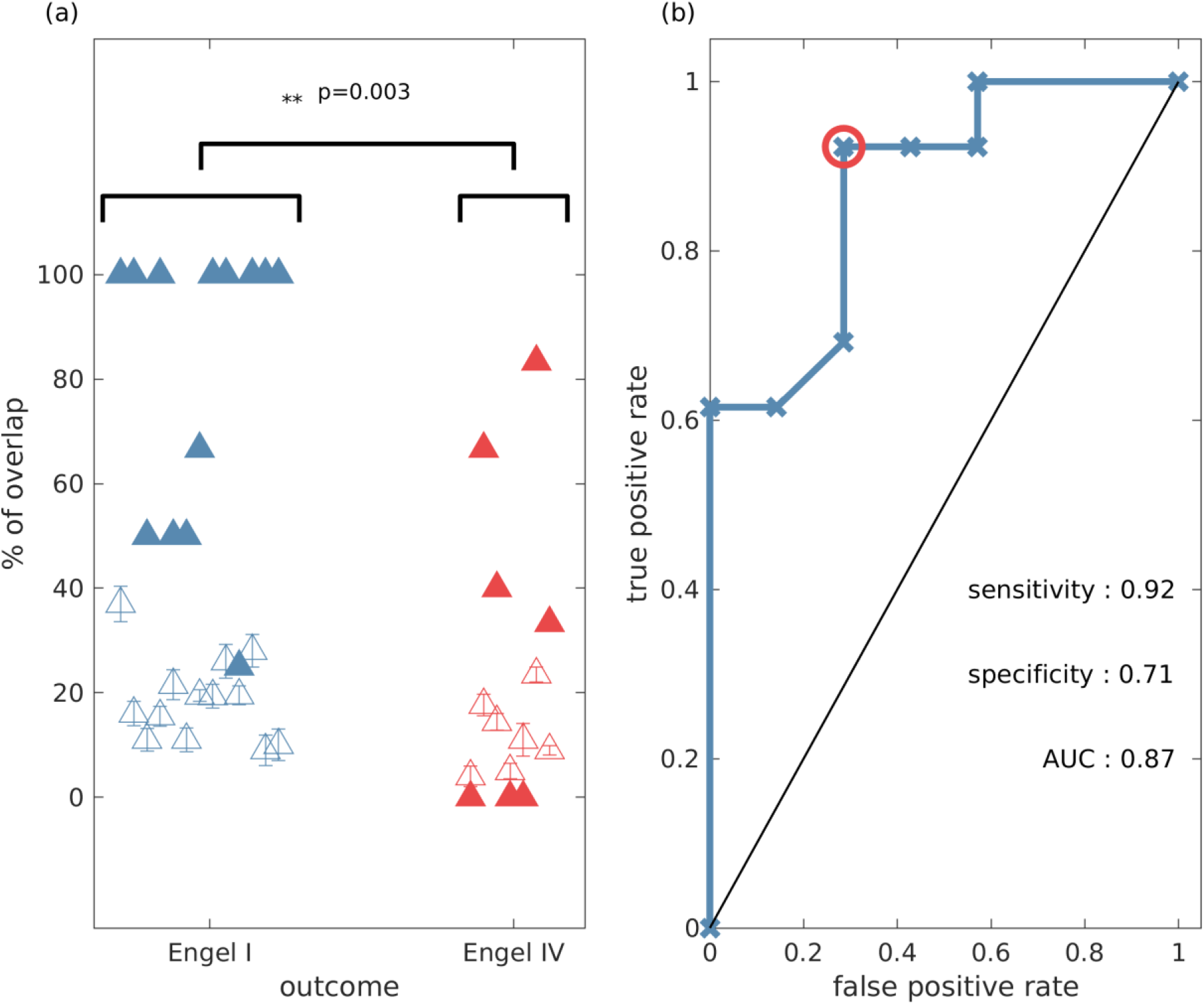
The percentage overlap between actual resections and model predictions is higher for patients with good outcome compared to those who had poor postsurgical outcome (one-sided Wilcoxon rank sum test *p* = 0.003). (a) Percentage overlap versus patient surgical outcome grouped by Engel class (filled triangles). The unfilled triangles correspond to an average overlap between actual resections and equal-size random resections (100 random samples, and the error bars denote the standard error). (b) Receiver operating characteristic (ROC) analysis for Engel I (seizure free patients) versus Engel IV (non-seizure free patients) using the percentage overlap as the classifier. Parameter setting for the genetic algorithm: population size equal 200 and number of generations equal 150.

For illustrative purposes, Fig 8 demonstrates the functional network of an Engel IV patient with two alternative optimal sets revealed by the genetic algorithm (Fig 8a and b). Here we demonstrate a further advantage of the genetic algorithm: it facilitates the avoidance of removing a certain node or nodes. This can be done by setting the objective function to a high value if the node(s) that should be avoided appear in a solution during the execution of the algorithm. Here, we avoid the selection of a highly ictogenic node (Fig 8c) and consequently the genetic algorithm substitutes it with another one. This constrained strategy of the genetic algorithm may be clinically valuable given that there may exist network nodes that cannot be removed due to their overlap with eloquent cortex, blood vessels or other anatomically indispensable areas.

**Fig 8.**
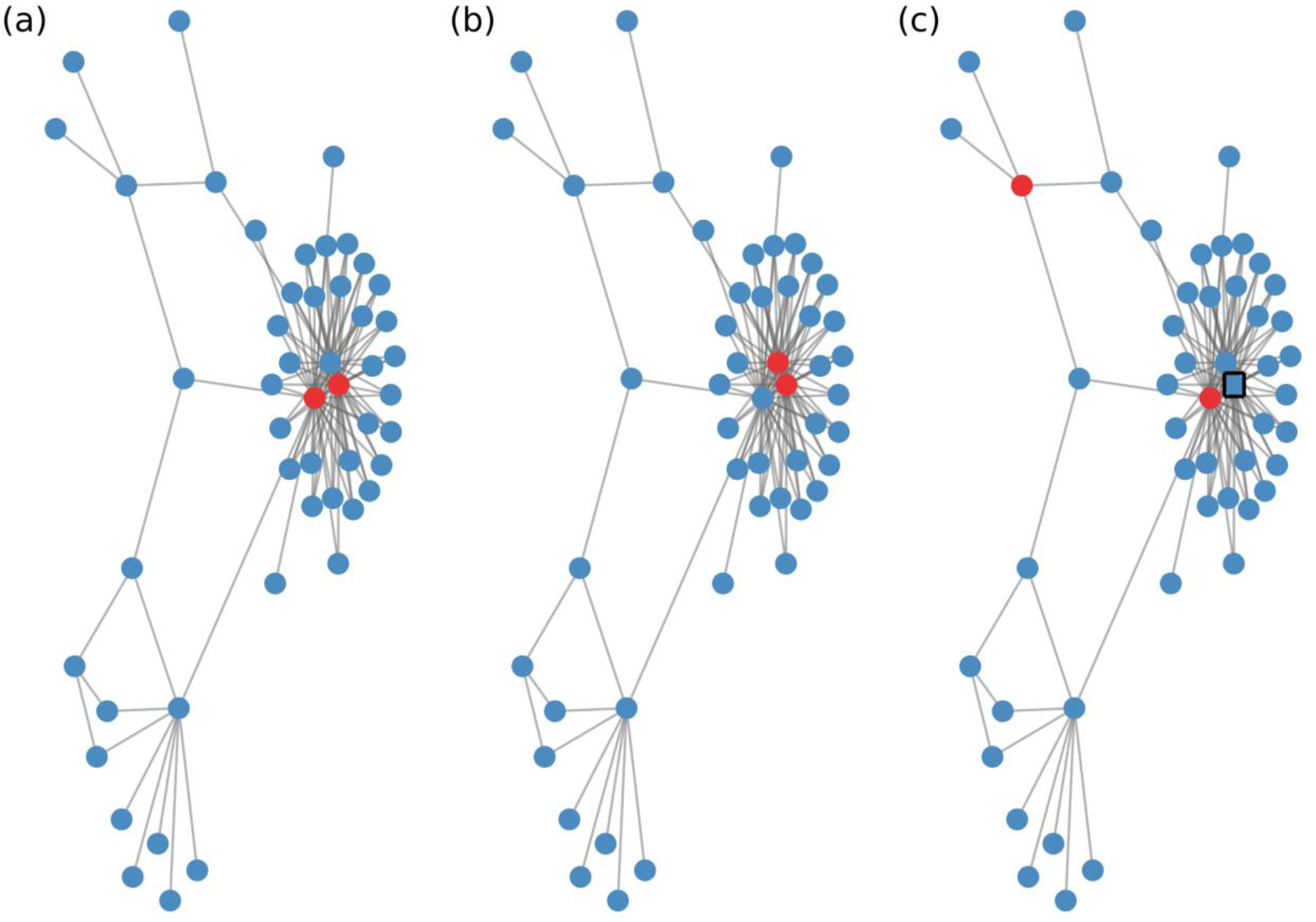
The genetic algorithm suggests multiple optimal sets and may further provide alternative sets under given restrictions. Each panel illustrates the functional network of an Engel IV patient as inferred from surrogate corrected mutual information. For illustrative reasons we only display the largest connected component of the network and connections characterized by correlations larger than 0.09. Nodes in red denote the optimal set as predicted by the genetic algorithm. Panels (a) and (b) exemplify two different (unconstrained) optimal sets. Panel (c) shows an additional alternative optimal set when constraining the genetic algorithm to avoid selecting the node within the black box. The actual clinical resection comprised four nodes which were not part of the displayed network component. Parameters are the same as in Fig 7.

## 4. Conclusions

In this study we used computational modelling and evolutionary optimisation to understand how sets of nodes within a network contribute to its seizure generating capability (i.e. its ictogenicity). To do this we used a quantity called Set Ictogenicity *(SI)*, which is a model-based quantification of the effect that removing a set of nodes has on the capability of a network to transition between healthy and epileptiform dynamics. We demonstrated that the way in which *SI* varies for different sets of nodes depends upon network topology. Whilst in exemplar 20-node random directed networks most sets of nodes have similar and large *SI*, in exemplar directed networks generated using the Barabasi-Albert model (i.e. “scale-free” networks), we observed a V-shaped distribution of *SI*. In the latter case, most sets do not yield a large reduction in ictogenicity when removed from the network. This difference is in part explained by the observed high correlation between *SI* and degree: since nodes in scale-free networks have higher degree variability, this leads to higher *SI* variability across sets of nodes. We further observed that *SI* is correlated with betweenness centrality. These results build upon our previous findings in Ref. (Lopes et al., 2017), where we analysed the correlation between graph theory measures and the Node Ictogenicity (which is the reduction in ictogenicity obtained by removing a single node). Our results add further evidence that targeting hubs, which would be sets of nodes with high average degree and betweenness centrality, is likely to be a good strategy for epilepsy surgery (Van Diessen et al., 2013, Zubler et al., 2015, Stam 2016, Lopes et al., 2017).

However, we demonstrated that although sets of nodes that induce maximum reduction in average degree or centrality typically have high *SI*, they are not necessarily sets with *the highest* SI. We therefore developed a global optimisation framework that can be deployed in general to find the optimal resection, given a network structure. Epilepsy surgery relies on the identification of brain regions that are responsible for the emergence of epileptiform activity, but wherever possible mitigating effects on normal brain function. We therefore studied two objectives: the *SI* and the size of the resection, using a multi-objective genetic algorithm. Genetic algorithms are optimisation methods based upon the process of evolution and natural selection. They have been widely applied in neuroscience (Whitley et al., 1990, Nevado-Holgado et al., 2012, Avramidis and Akman 2017, Wang et al., 2018) and it has been shown that they are a valuable tool to solve combinatorial type computational problems (Hromkovic et al 2010, Luque et al., 2011). Here, we showed that in small networks our genetic algorithm was able to find minimal sets with highest ictogenicity. In larger networks, we compared the genetic algorithm with other heuristic approaches and demonstrated that the genetic algorithm was always at least as good or better than these heuristics at finding the sets with highest ictogenicity.

Brain networks may be studied at different spatial and temporal scales using different data modalities (Bassett et al., 2017). In the context of epilepsy surgery, most studies have focused on large-scale brain networks inferred from iEEG (Khambhati et al., 2015, Bartolomei et al., 2017) and MRI (Proix et al., 2017, Taylor et al., 2018). Network topology has been shown to evolve during seizures (Kramer et al., 2010, Lehnertz et al., 2014), with structures changing from random to more regular in seizures and back to more random after seizure termination (Schindler et al., 2008). Furthermore, it has been shown that hubs may play a crucial role in the generation of seizures (Wilke et al., 2011, Varotto et al., 2012, Lopes et al., 2017). That is why here we studied both random and scale-free networks to build understanding about the epileptic brain. We found that the *SI* distribution varies in networks with different topologies and is more heterogeneous in directed scale-free networks compared to directed random networks. The framework we introduced is flexible and can be applied to smaller spatial scales, e.g. neuronal networks or smaller sized regions of interest in whole brain models (Stead et al., 2010, Smith et al., 2016). At the smaller scale, these methods could shed light on, for example, why hubs of granule cells in the dentate gyrus are responsible for the emergence of seizures after brain injury (Morgan et al., 2008). Use of these methods combined with experimental testing of perturbations, for example using optogenetics (Chow et al., 2013), could open up new avenues for targeted treatment for seizures (McGovern et al., 2016).

We demonstrated the potential applicability of the genetic algorithm by applying it to functional brain networks derived from iEEG recordings from 20 patients who had undergone epilepsy surgery. We found that the model-derived optimal set had larger overlap with actual resections in the case of patients who were ultimately seizure free. This is in line with previous studies that have used computational models and heuristic approaches, based on properties of individual nodes, to test potential alternative resections (Goodfellow et al., 2016, Sinha et al., 2017, Lopes et al., 2017). Furthermore, our framework achieved a classification performance comparable to recent studies that used machine learning and quantitative EEG methods (Müller et al 2018). Depending on the way in which ictogenic networks are constructed, it is possible that in epilepsy surgery multiple nodes from the brain network are removed. We therefore here aimed to identify the indispensable brain region for seizure generation by considering the ictogenicity of sets of nodes. A particular advantage of the genetic algorithm that we demonstrated is that it naturally suggests multiple optimal sets that suppress the epileptiform activity of the network, due to the random nature of the search. These are alternative sets that give rise to optimal *SI* (see Figs 8a and b). Furthermore, there are natural ways to implement constraints, for example on nodes that should not be removed because they are essential for healthy brain function (Fig 8c).

There are a number of caveats to the approaches we outlined. We validated the approach by measuring the overlap of the optimal set with actual resections, taking into account postoperative seizure freedom. Although the results of Fig 7 give confidence that the collection of nodes identified in the analysis were in fact ictogenic, it does not provide validation that the entire set itself is the optimal resection. In order to test this further, we propose to work with experimental systems, whereby alternative resections can be performed (Sheybani et al., 2018). We note that a large overlap between the model suggestion and the actual resected tissue was found in 2 cases in which outcome was Engel IV. I In addition, in one Engel class I patient the overlap was small, see Fig 7a. Our approach assumes the existence of an ictogenic network and it has been shown that even focal epilepsies may involve in the seizure generation mechanism widespread brain regions (Spencer 2002, Richardson 2012, Bartolomei et al., 2017, Besson et al., 2017). However, the electrodes are implanted in a designated brain region and the functional network that is inferred from them might not reflect the ictogenic network. Therefore, the initial placement of iEEG electrodes may be key here, and recent work has aimed to use modelling to uncover cases for which an alternative implantation scheme may be required (Lopes et al., 2017). Future work should also aim to aid clinicians with regards to electrode implantation and to integrate different data modalities so that predictions may be more robust. We further note that although *SI* can be used to compare different resections, their actual values can be difficult to interpret. Linking *BNI* and *SI* to the rate of occurrence of epileptiform discharges in humans and experimental models will be an important avenue for future work that can aid the refinement of model predictions. We also note that the genetic algorithm is computationally more expensive than the other heuristics (see Supplementary material, Text S3), however, it may be further optimized in the future by making use of parallelisation and GPUs (Avramidis et al., 2017), for example.

In conclusion we presented a computational approach that quantifies the contribution of brain regions to seizure generation. Our approach enhances the understanding of how perturbations in brain networks may lead to seizure freedom. It allows multiple surgical strategies to be tested *in silico* in order to find the optimal set that reduces the network ictogenicity. In addition, the genetic algorithm that we deployed finds the optimal trade-off between the size of the resected tissue and the reduction in network ictogenicity. Our results show promise that the computational approaches introduced herein have the potential to be incorporated into surgery decision pipelines in practice.

## Acknowledgments

We would like to acknowledge Khulood Alyahya, Kevin Doherty, and Leandro Junges for fruitful discussions and comments. MG, M.A.L and MR gratefully acknowledge funding from the Medical Research Council [grant number MR/K013998/1]. MG and MR received financial support from the EPSRC [grant number EP/N014391/1]. MG and PL received financial support from the EPSRC [grant number EP/P021417/1]. The contribution of MG was further generously supported by a Wellcome Trust Institutional Strategic Support Award [grant number WT105618MA]. MR is also funded by the NIHR Biomedical Research Centre at South London and Maudsley NHS Trust and King’s College London; and by the Medical Research Council Centre for Neurodevelopmental Disorders at King’s College London [MR/N026063/1]. OEA would like to acknowledge the financial support of the EPSRC [grant numbers EP/N017846/1 and EP/N014391/1]. CR and MM received support from the Swiss League Against Epilepsy. El. A acknowledges funding from EPSRC Tier-2 capital grant EP/P020259/1. Eu. A is funded by the European Union’s Horizon 2020 research and innovation programme under the Marie Sklodowska-Curie grant agreement no. 75088.

## Abbreviations

iEEG: intracranial electroencephalographic recordings
EZ: epileptogenic zone
*BNI*: Brain Network Ictogenicity
*NI*: Node Ictogenicity
*SI*: Set Ictogenicity

