## Supplementary material for "Quantification and selection of ictogenic zones in epilepsy surgery": Supplemetary material

### Text S1: Derivation of functional networks from iEEG recordings

In order to derive the functional networks from the iEEG recordings we followed the methods described in Rummel et. al (Rummel et al., 2013, Rummel et al., 2015). We first divided the signals in segments of 8 seconds and generated 10 sets of multivariate iterative amplitude adjusted Fourier transform (IAAFT) surrogates for each segment independently. Afterwards, each segment from both the original and surrogate signals was divided in 10 subsegments of 2 seconds each with minimal overlap. Surrogated corrected mutual information matrices were then calculated by computing mutual information between pairs of signals both from the 10 subsegments from the original signals and the 100 subsegments from the surrogates. The correction was performed by assessing the statistical significance of the mutual information as obtained from the original signals compared to the surrogates using a Mann–Whitney U test. We applied this process for each segment, and therefore we obtained time-varying functional connectivity networks. For our analysis we used the segments corresponding to the first half of seizures, because it has been shown (Rummel et al., 2015) that functional networks derived from these epochs correlate well with epileptogenic tissue.

### Text S2: $BNI$ and $SI$ calculation

The $SI$ calculation is preceded by finding the scaling factor $K$ such that $BNI^0 = 0.5$. We used a root-finding algorithm for this. Note that the $K$ value and all other parameters are then kept constant for the $SI$ calculation. Regarding the $BNI$ calculation, we first define whether a node is spiking by applying a threshold to the signal of each node (more precisely, the signal is $(1/2)(1 - \cos(\theta_i(t) - \theta_i^s(t))$ and we defined spiking activity as any signal larger than 0.9). Spikes less than 2400 time steps apart were considered as belonging to the same spiking epoch. $BNI$ was then the average fraction of time that each node spent in spiking activity during a sufficiently long reference time ($4 \times 10^6$ time steps). We computed five noise runs and then calculated the average $BNI$ value. Note that in the case of the recurrent ordering approach, after each node removal, the reference $BNI^0$ had to be recomputed. However, in some cases, after the removal of a number of nodes, $BNI$ could not reach 1. To account for this, the reference $BNI^0$ was set to half of the maximum available $BNI$ at each iteration of this strategy.

#### Text S3. Comparison of the computational time of all resection strategies

To compare the computational time that each strategy requires, let us consider a network with $N$ nodes and, as unit of computational cost, the time that is required for the $SI$ calculation of a set of nodes regardless of its size. Then, the ground truth strategy requires $2^N - 1$ units, the simple ordering $N$, the recurrent ordering $\sum_{i=0}^{N-1}(N - i)$, and the genetic algorithm requires the product (number of generations) x (population size) x (number of independent iterations of the Algorithm) units to identify the most ictogenic set. For typical network sizes $N < 150$, the computational cost of the genetic algorithm is higher than both simple and recurrent ordering but much cheaper than the ground truth.

### Supplementary Figures

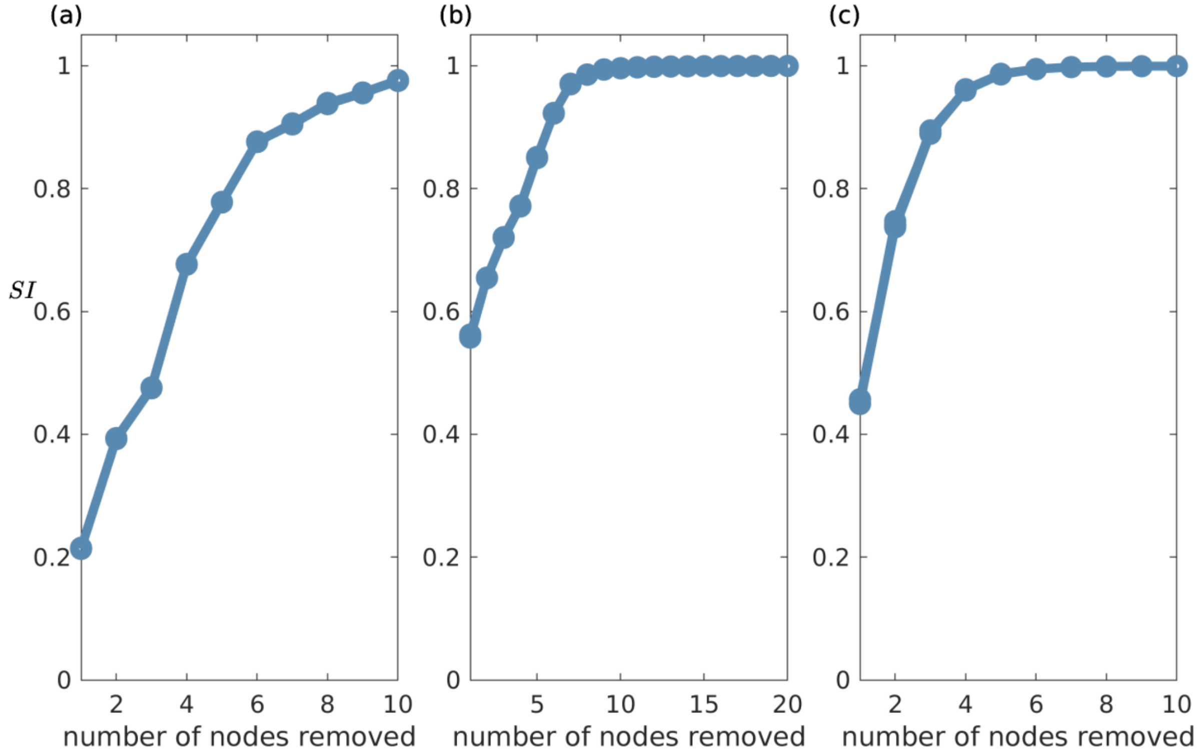


**Fig S1.** Set Ictogenicity values of the most ictogenic sets as detected by the genetic algorithm across different resection sizes. Panel (a) and (b) show $SI$ in 20- and 40-node directed scale-free networks, respectively. The $SI$ in panel (c) was computed in a functional network derived from iEEG recordings. Each panel shows eight overlapping curves that correspond to eight independent runs of the genetic algorithm. The concordance between different runs provides confidence on the observed solutions.

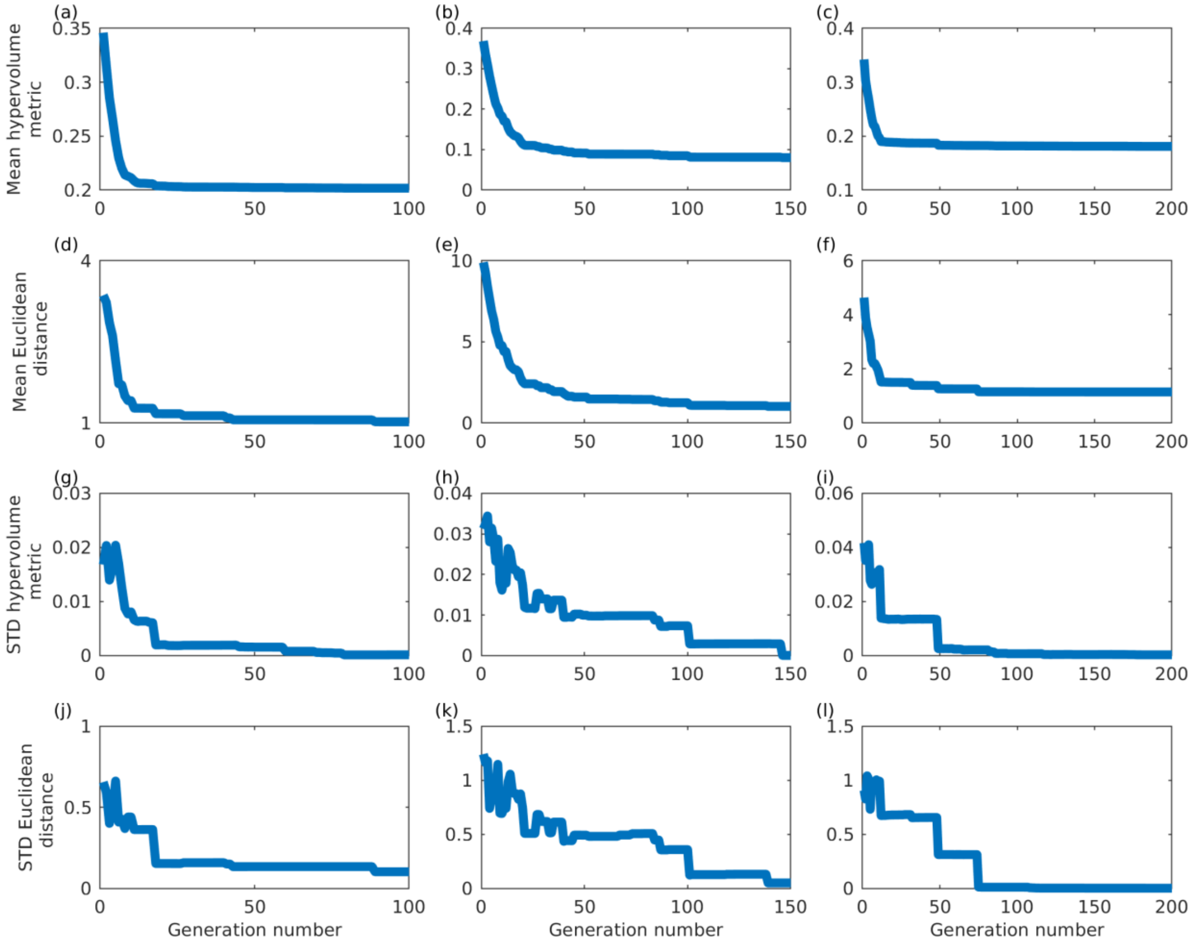


**Fig S2.** Convergence metrics versus the generation number. The choice of convergence metrics was discussed elsewhere [Avramidis and Akman 2017 and references therein]. Each row shows a different convergence metric: mean hypervolume metric, mean Euclidean distance, STD hypervolume metric, and STD Euclidean distance. Panels (a, d, g, j), (b, e, h, k) and (c, f, i, l) correspond to the 20-node, 40-node and the patient network shown in Fig S1 respectively.

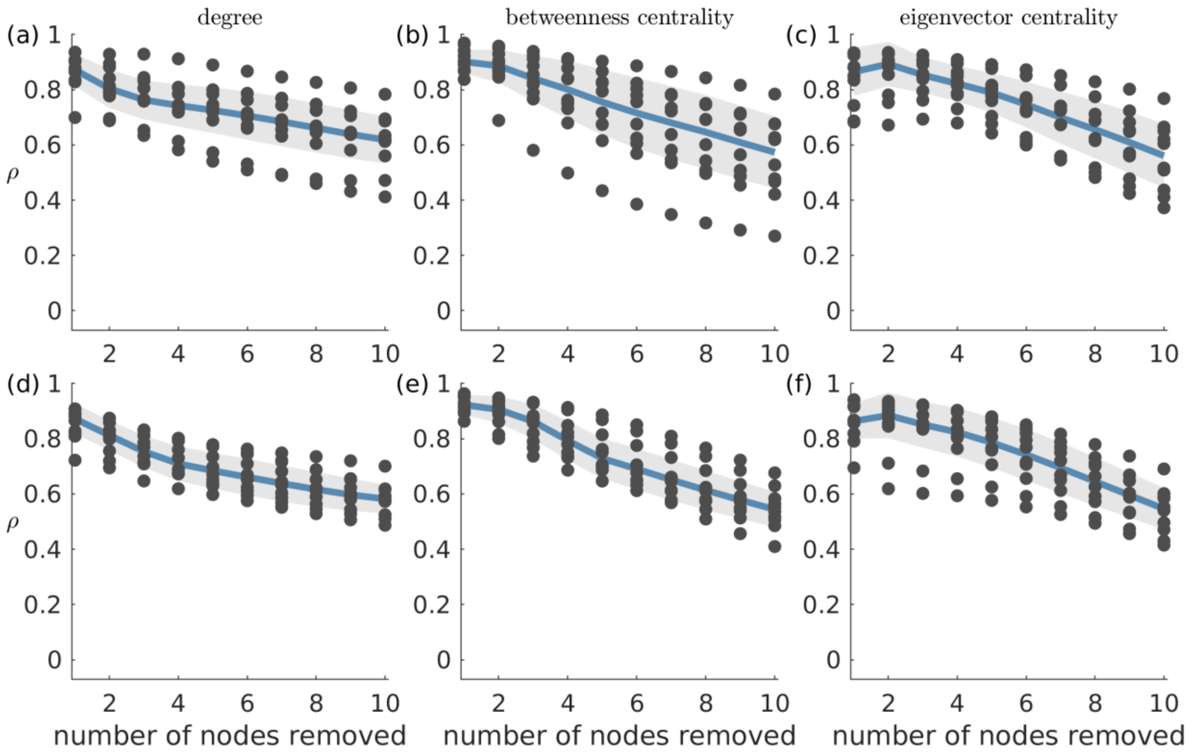


**Fig S3**. Absolute Spearman's correlation ($\rho$) between $SI$ and average graph theory measures of the nodes in removed sets. This figure shows the same as Fig. 3 except that here we consider undirected networks. Panels (a)-(c) correspond to scale-free networks while panels (d)-(f) stand for random networks. Each column shows $\rho$ between $SI$ and a different network measure: (a) and (d) average degree; (b) and (e) average betweenness centrality; and (c) and (f) average eigenvector centrality of removed nodes. Ten network realizations were considered per network topology, hence the 10 dots for each resection size (i.e. number of nodes removed). The blue line represents the median across the network realizations and the shaded area displays the median absolute deviation. The parameters were the same as in Fig 2. The mean degree of all the considered networks was two.

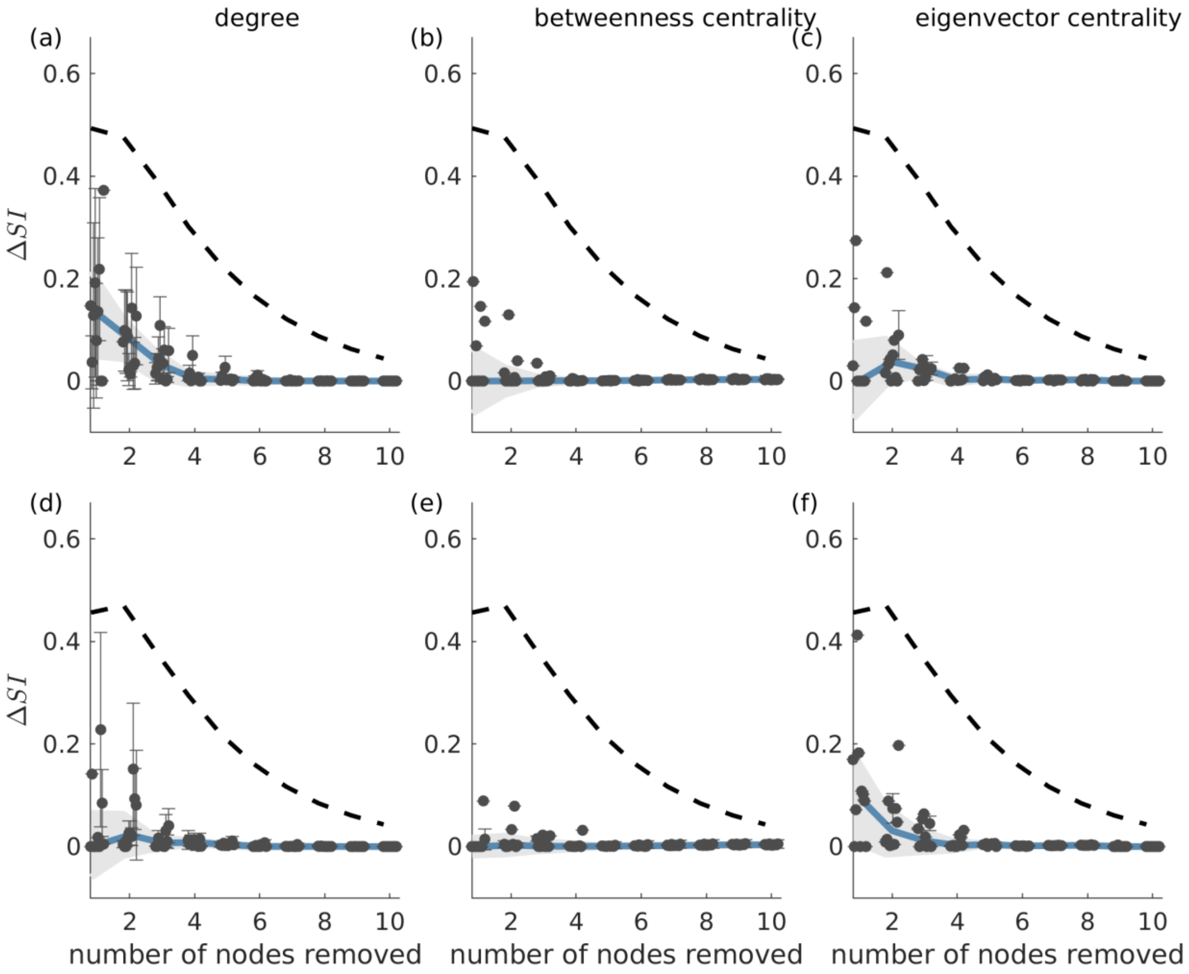


**Fig S4.** Difference between the $SI$ value of the most ictogenic set as identified from the ground truth and the average $SI$ of the sets which caused a maximal reduction in average degree (panels (a) and (d)), average betweenness centrality (panels (b) and (e)) and average eigenvector centrality (panels (c) and (f)) i.e. $\Delta SI$, as a function of resection sizes. This figure shows the same as Fig. 4 except that here we consider undirected networks. Error bars denote the standard deviation of the $\Delta SI$ values across the different sets that yield the maximal reduction in average degree or betweenness centrality when removed. The blue curve describes the median of the $\Delta SI$ values across 10 network realizations (black dots) and the shaded area their median absolute deviation (the dots are slightly shifted in the x-axis for better visualization). Panels (a)-(c) correspond to scale-free directed networks, whilst panels (d)-(f) to random undirected networks. The dashed line denotes the average $\Delta SI$ between the $SI$ of the ground truth most ictogenic set and the $SI$ of all other possible sets (also averaged over the 10 network realizations). Parameters are the same as in Fig 2.

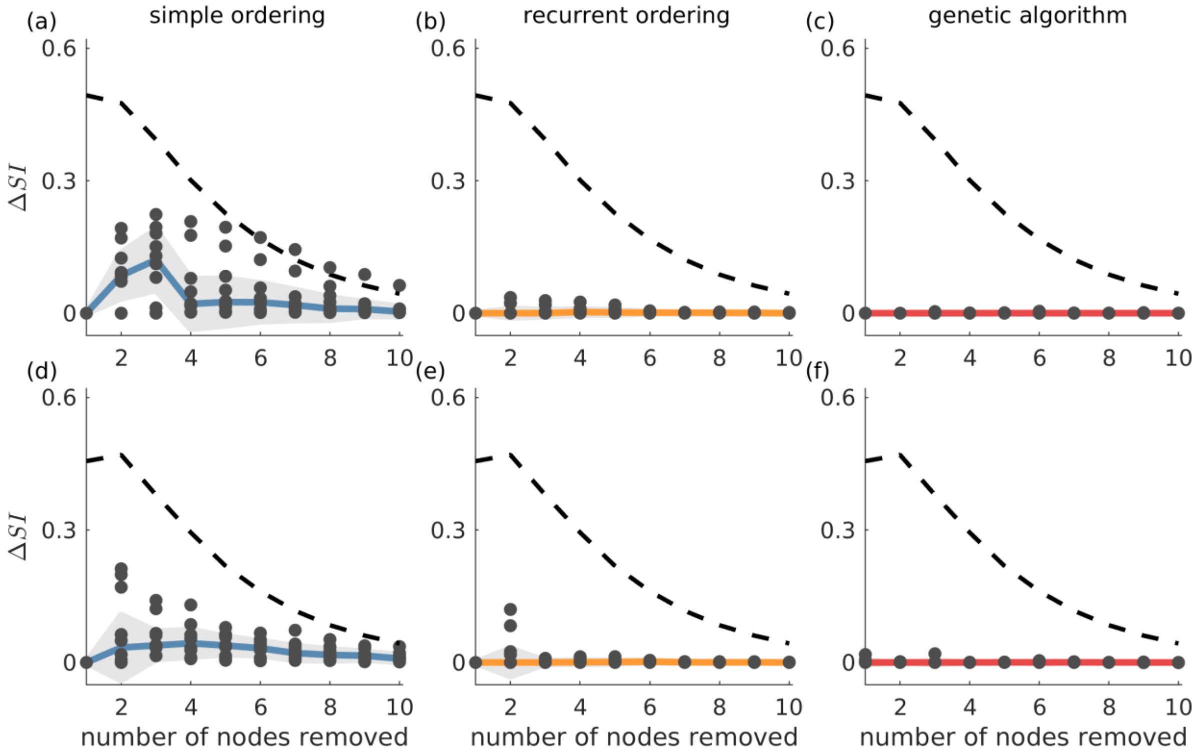


**Fig S5.** $\Delta SI$ versus the number of resected nodes. The ground truth is compared with simple ordering (first column), recurrent ordering (second column) and the genetic algorithm (third column). This figure shows the same as Fig 5 except that here we consider undirected networks. The first and second rows show the $\Delta SI$ between the $SI$ value of the ground truth most ictogenic set and the one identified by each strategy in scale-free and random networks, respectively. The dashed lines describe the average $SI$ (or $\Delta SI$) of all possible combinations of nodes for each resection size and serve as a reference for comparison. The dots correspond to $\Delta SI$ values obtained for 10 different network realizations, the solid lines depict the median of $\Delta SI$ values, and the shaded area represents the median absolute deviation. Parameters were the same as in Fig 2. Additionally, the genetic algorithm was run with 100 generations and population size 200. The mean degree in all networks was two.

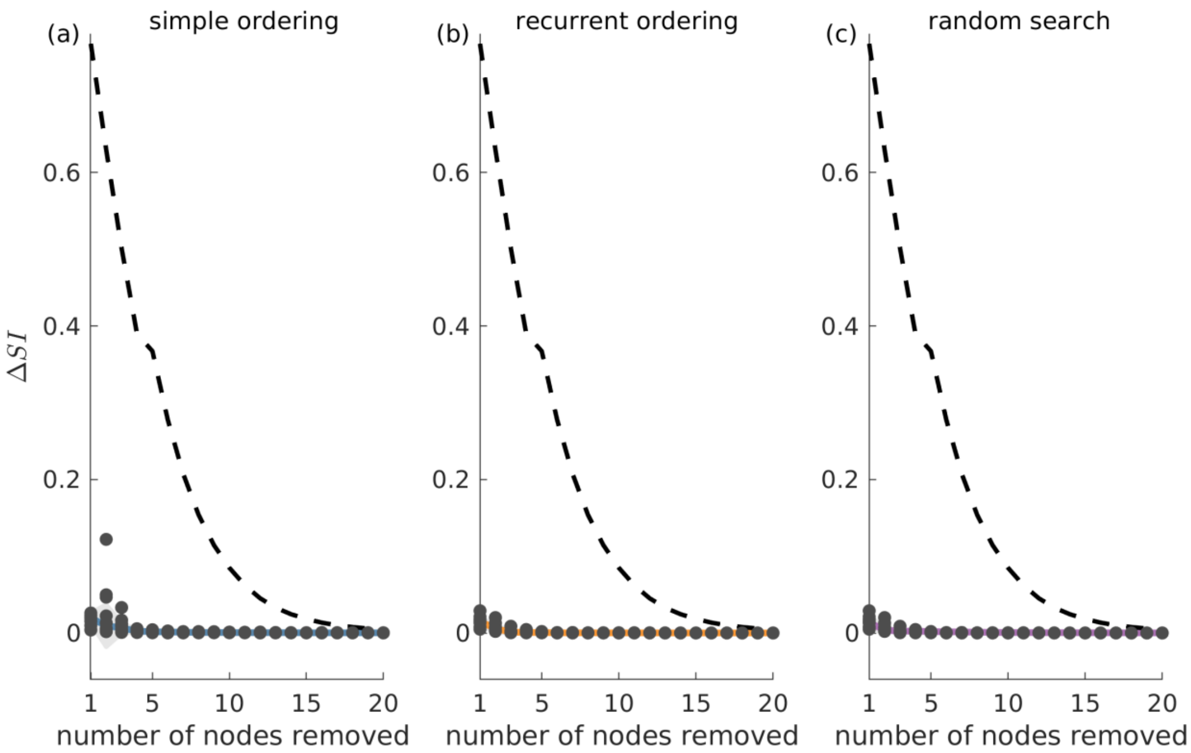


**Fig S6.** All resections strategies find similar solutions in 40-node scale-free undirected networks. $\Delta SI$ denotes the difference between the $SI$ value of the optimal set as detected by the genetic algorithm and the $SI$ solution found by (a) simple ordering, (b) recurrent ordering, and (c) random search across different resection sizes. The solid lines depict the median of the dots which correspond to the $\Delta SI$ values across 10 network runs, whilst the shaded area illustrates the median absolute deviation. The dashed line represents the difference between the optimal set as detected from the genetic algorithm and 20,000 random sets for each resection size (for resections up to three nodes, we considered all possible sets, since they were less than 20,000). Parameters: mean degree equal 4; population size 200, number of generations equal 150.

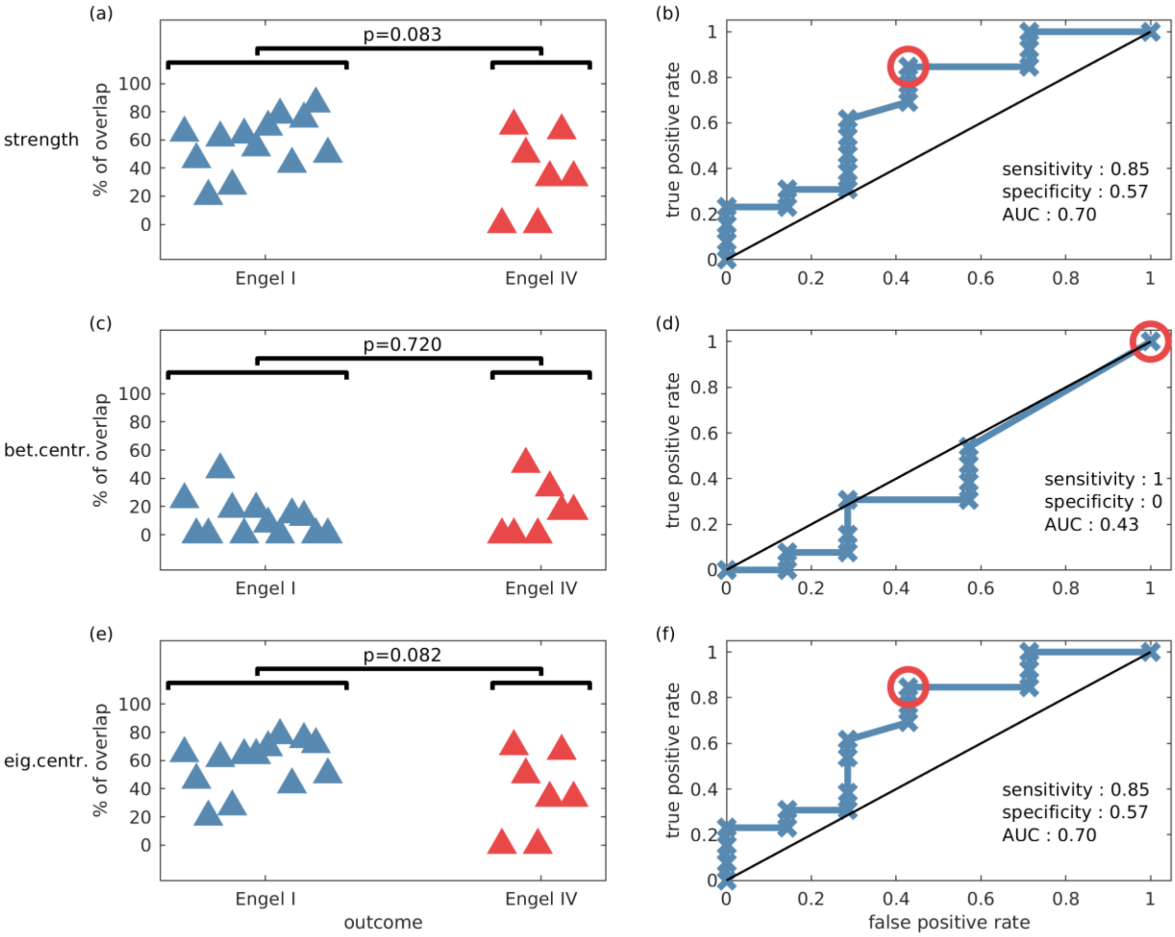


**Fig S7.** The percentage overlap between actual resections and predictions based on graph theory measures (with predicted set size equal to the size of the actual resected set). Each row is comparable to Fig 7, i.e. the panel on the left shows the percentage overlap versus patient surgical outcome grouped by Engel class (where one-sided Wilcoxon rank sum tests were used), and the panel on the right is the receiver operating characteristic (ROC) analysis for Engel I (seizure free patients) versus Engel IV (non-seizure free patients) using the percentage overlap as the classifier. Each row corresponds to a different graph theory measure: (a-b) node strength (i.e. the sum of connection weights), (c-d) betweenness centrality, and (e-f) eigenvector centrality. All these measures are incapable of achieving performances comparable to the predictions based on *SI* (see Fig. 7).

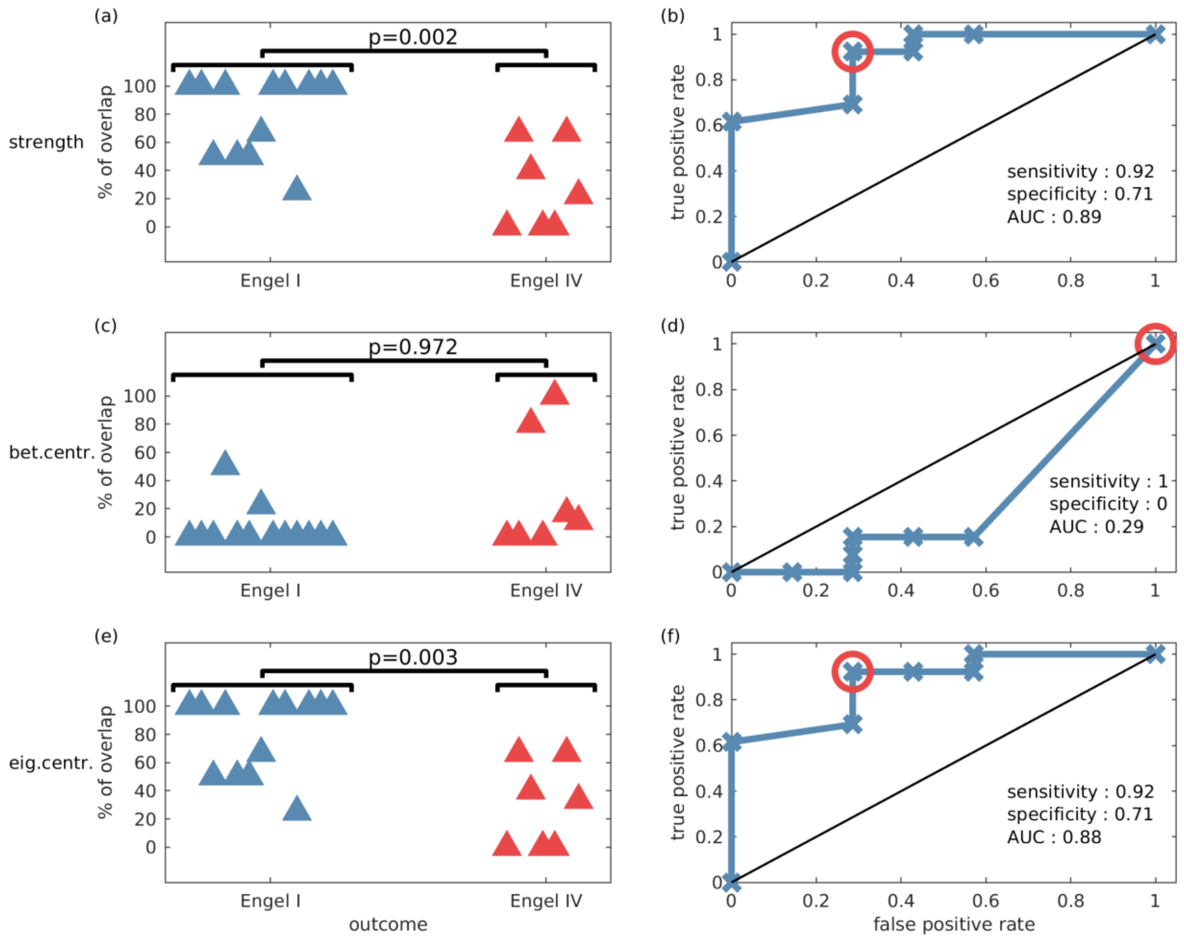


**Fig S8.** The percentage overlap between actual resections and predictions based on graph theory measures (with predicted set sizes equal to the size of predicted sets using *SI* in Fig. 7). Each row is comparable to Fig 7, i.e. the panel on the left shows the percentage overlap versus patient surgical outcome grouped by Engel class (where one-sided Wilcoxon rank sum tests were used), and the panel on the right is the receiver operating characteristic (ROC) analysis for Engel I (seizure free patients) versus Engel IV (non-seizure free patients) using the percentage overlap as the classifier. Each row corresponds to a different graph theory measure: (a-b) node strength (i.e. the sum of connection weights), (c-d) betweenness centrality, and (e-f) eigenvector centrality. The strength and eigenvector centrality are capable of achieving performances comparable to the predictions based on SI (see Fig. 7). Note, however that the predicted set size was informed by the *SI* calculation.
